# Probability of stealth multiplets in sample-multiplexing for droplet-based single-cell analysis

**DOI:** 10.1101/2023.12.22.573050

**Authors:** Fumio Nakaki, James Sharpe

**Affiliations:** European Molecular Biology Laboratory (EMBL) Barcelona, Carrer Dr. Aiguader 88, 08003, Barcelona, Spain; Institució Catalana de Recerca i Estudis Avançats (ICREA), Passeig Lluís Companys 23, 08010, Barcelona, Spain

**Keywords:** single-cell analysis, scRNA-seq, sample multiplexing, Poisson distribution, stealth multiplet

## Abstract

**Background:** Recent advances in sample-multiplexing droplet-based single-cell RNA sequencing (mx-scRNA-seq) enable us to evaluate large numbers of different samples or experiments simultaneously by reducing the occurrence of undetectable multiplets, that is, the droplets that capture multiple cells. However, the probability of potential multiplets in mx-scRNA-seq is yet to be quantitatively examined.

**Results:** We developed a simple theoretical model to predict four classes of possible multiplets in mx-scRNA-seq: Homogeneous stealth, partial stealth, multilabelled, and unlabelled. We estimated the probability of each class and have found that the partial tsealth multiplet, which has been overlooked, may substantially affect the whole data when the labelling process is suboptimal. Next, we illustrated their presence in actual mx-scRNA-seq datasets when the sample labelling was suboptimal. In addition, we found that choosing a suitable sample-demultiplexing algorithm was critical to circumvent the partial stealth multiplets.

**Conclusion:** Our results show the necessity of optimising the labelling procedure and offer a theoretical basis to estimate the probability of each type of multiplets to ensure the integrity of mx-scRNA-seq.

## Background

Characterising every cell in a given biological system is one of the major goals of the analytic approach in biology. Recent advances in single-cell transcriptome analysis have greatly expanded the number of genes and cells for analysis, allowing to characterise a massive number of individual cells simultaneously, depicting a representative transcriptome landscape of the biological system of interest and providing us with a new insight into complex biological systems [1–7].

The microfluidic-based platforms are among the technological platforms for single-cell RNA sequencing [1, 2, 6] (hereafter, we use the term “scRNA-seq” specifically for the microfluidic-based random cell capture method unless otherwise denoted). They are commercially available and widely accessible without expertise in single-cell biology [8]. This technology utilises a droplet generated in a microfluidic circuit that contains a bead with a unique nucleotide barcode. When a cell suspension is loaded, each cell is partitioned and isolated in a single droplet, and transcripts are directly captured and labelled with nucleotide barcodes attached to a bead. Although it can theoretically label as many cells as the variation of the barcode sequences allows, one of the critical limiting factors in practice is the presence of multiplets, droplets having more than one cell [1, 2, 6]. The more cells are loaded into the microfluidic device, the greater the risk of multiplets, which threatens the integrity of scRNA-seq by undermining the basis that each transcriptome comes from a single cell. Controlling the frequency of multiplets is, therefore, one of the critical quality control steps in scRNA-seq. One direction to tackle this issue is to develop an algorithm to detect the multiplets from their transcriptome pattern. Several multiplet removers *in silico* have been proposed [9–12]. However, this approach is not an optimal solution because it has yet to distinguish the “homotypic” multiplets that consist of cells having similar transcriptome profiles [9–12]. Also, it utilises the transcriptome data in question for predicting the multiplets that should be removed. In other words, it may pose a risk that the subsequent transcriptome analysis falls into circular reasoning. Another strategy is to facilitate the multiplet detection by sample labelling [13–18] or inherent genetic variation [19, 20] before loading onto the microfluidic device. These additional sample labels reveal the multiplets that appear as cell-droplets with multiple sample labels, which, in turn, raises the ceiling of cell number per single run by mitigating the risk of undetectable multiplets [14, 21]. Moreover, it is advantageous to accurately compare multiple samples by removing the inevitable technical batch effect when multiple samples are loaded individually, broadening the application of scRNA-seq [18].

However, even though the potential multiplets can be efficiently removed by analysing the sample barcodes of each droplet that contains at least one cell (hereafter “cell-droplet”) in mx-scRNA-seq, there is still a concern that the cells classified as monolabelled cells are not always true singlets. For example, a multiplet comprising multiple cells with the same sample barcode cannot be detected by barcode demultiplexing [12, 14, 21, 22]. Another potential form of hidden multiplets is a combination of labelled and unlabelled cells, which has not been examined so far. Estimating and controlling these hidden multiplets among apparently monolabelled cell-droplets - we call them “stealth multiplets” hereafter - is essential for experimental design and data analysis in the mx-scRNA-seq. Still, there has been no quantitative assessment of the probability of these different types of stealth multiplets.

In this article, we propose four categories of multiplets in mx-scRNA-seq from the viewpoint of sample barcoding: (1) the multiplet of all cells with the same sample barcode and detected as monolabelled (homogeneous stealth), (2) the multiplet that is a combination of a labelled cell(s) of single barcode and an unlabelled cell(s) and detected as monolabelled (partial stealth), (3) the multiplet contains the cells with different sample barcodes (multilabelled), and (4) the multiplet consisting of unlabelled cells (unlabelled). While the multilabelled multiplet and unlabelled multiplet can be eliminated easily, the homogeneous and partial stealth multiplets are undetectable at the demultiplexing step. We estimate the probabilities of each type of multiplet based on a Poisson distribution model and examine their behaviour depending on the parameters, including a cell loading rate, labelling efficiency, and sample numbers to be multiplexed. Interestingly, the partial stealth multiplet cannot be controlled by increasing the number of multiplexed samples, but the labelling efficiency is a critical parameter for their occurrence. Moreover, since the labelling efficiency is affected not only by the sample labelling procedure *in vitro* but also by the demultiplexing methods *in silico*, we suggest testing multiple demultiplexing algorithms to ensure accurate downstream analysis by minimising the risk of the stealth multiplet.

## Results

### Theoretical probability of stealth multiplets occurred in monolabelled droplets

The probability of a cell-droplet that contains a given number, *k*, of cells can be assumed to follow the Poisson distribution [6, 10, 22, 23],

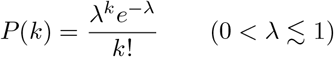

where *λ* is the average count of the event, representing the average cell numbers per droplet, which must be 0 *< λ* ≲ 1 in practice because cells should be loaded with low concentration to generate multiplets as little as possible, which also means *λ* is closely correlated with cell loading rates in practice. With this model, the probability of empty droplets, singlets, doublets, and *k*-ets (with or without beads) is *P* (0), *P* (1), *P* (2), and *P* (*k*), respectively (See **Supplementary information** for further discussion).

In addition, to examine the composition of cell-droplets in mx-scRNA-seq, we need to consider the sample labelling (or cell hashing or cell tagging) status. Genetic variation (e.g. single-nucleotide polymorphisms) can be a stable cell label [19, 20], but is not readily applicable to every experimental setting. On the other hand, inducing external labels such as oligonucleotide-conjugated antibodies [13, 15], lipid-conjugated oligonucleotide [14], click chemistry [17], or nucleotide transfection [16], is more versatile. However, because of the inherent randomness in these labelling reaction processes, there is a fraction of poorly labelled cells even in a well-labelled sample, and these cells are potentially classified as unlabelled cells on demultiplexing after sequencing.

Here, we first introduce a conceptual “singleplex” experiment in which only one label is used. In this hypothetical setting, cell droplets can be classified by how many cells are encapsulated and whether they are labelled or not (**Figure 1A**). For simplicity, hereafter, we divide the whole population from a given sample into label-positive cells (labelled cells) and label-negative cells (unlabelled cells) and define the “labelling efficiency”, *a* (0 *< a <* 1), as a ratio of labelled cells among the whole population.

**Fig. 1:**
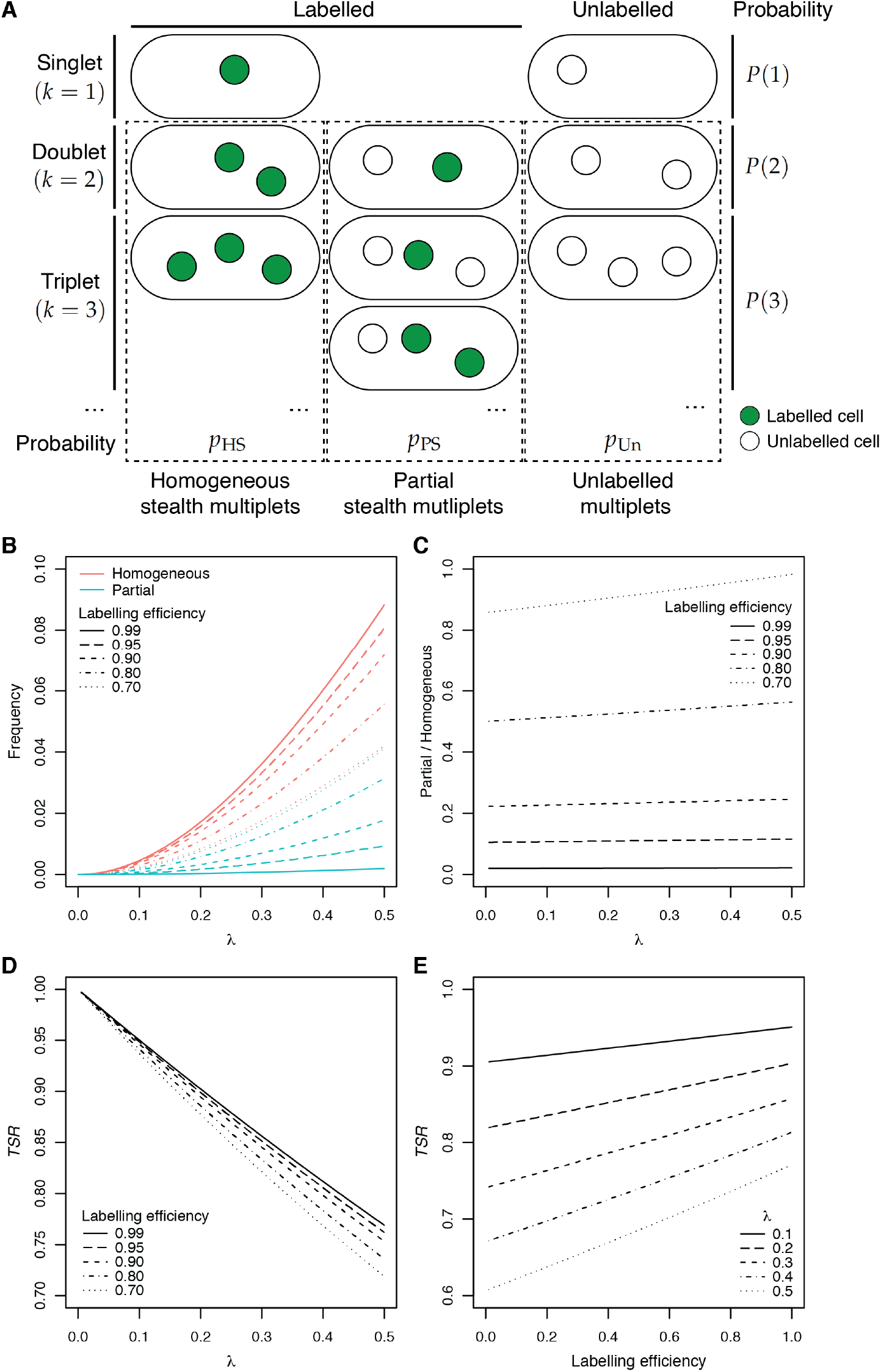
Stealth multiplets as multiplets among monolabelled cells in an conceptual singleplex experiment. (A) Schematic illustration of the combinations of cells in a single droplet with single-barcode labelling. Labelled cells are shown in green and unlabelled in white. The probability of the whole singlet, doublet, triplet,…, can be expressed as *P(1), P(2), P(3)*,…, respectively. Also, the parts of probability for the homogeneous stealth, partial stealth, and unlabelled multiplets are expressed as *p*_HS_, *p*_PS_, and *p*_Un_, respectively. (B) The frequency of the homogeneous and partial stealth multiplet among all cell-droplets under various *λ* and labelling efficiencies. (C) The ratio of homogeneous and partial stealth multiplet is plotted against labelling efficiencies under various *λ*. Note that the ratio is close to 1 when the labelling efficiency is low. (D) *TSR* (true singlets among the monolabelled cell-droplets) is plotted against *λ* under various labelling efficiencies. (E) *TSR* is plotted against labelling efficiency under various *λ*.

The sample labelling process is upstream of the droplet formation, and, therefore, the labelling efficiency is independent of *λ*, the frequency of cell-droplet formation. Given that the frequency of the labelled and unlabelled cells are *a* and (1 *− a*) by definition, the probability of labelled singlets is *aP* (1). However, the probability of the monolabelled cell-droplet is not *aP* (1) because stealth multiplets are possible (**Figure 1A**).

In this scenario, the probability of the homogeneous stealth multiplet *p*_HS_ and the unlabeled multiplet *p*_Un_ can be expressed by infinite series as follows:

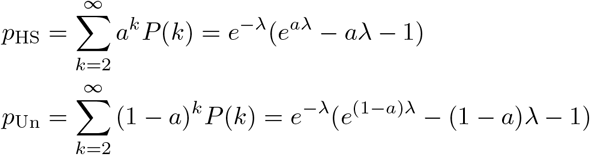

where *P* (*k*) is a Poisson distribution function. From these equations, the probability of the partial stealth multiplets *p*_PS_ is,

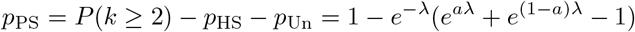

The *p*_HS_ and *p*_PS_ under various *a* and *λ* are shown in **Figure 1B**. The homogeneous stealth multiplet is dominant for very high labelling efficiencies (*≥*99%), and few partial stealth multiplets are present, as expected. However, the ratio of homogeneous to partial stealth multiplets (**Figure 1C**) is close to 1 when the labelling efficiency is 70%.

Next, we estimate the true singlet ratio (*TSR*), the fraction of true singlets among monolabelled droplets. The probability of the true singlet is *aP* (1). Therefore,

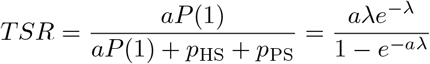

As *λ* increases, the overall multiplet ratio increases, and the *TSR* decreases irrespective of the labelling efficiency (**Figure 1D**). Note that the *TSR* is 90%-95% when *λ* is 0.1-0.2, which is estimated to correspond to the target cell number 6,000-12,000 with the Chromium Controller (10X Genomics) (Detailed discussion on estimated *λ* in the Chromium Controller is described in **Supplementary information**). This low *TSR* is because barcode demultiplexing cannot eliminate multiplets in this hypothetical singleplex experiment. The relation between *TSR* and labelling efficiency for various *λ* is shown in **Figure 1E**. The slope becomes steep as *λ* increases, suggesting that the labelling optimisation becomes important, particularly when a large number of cells are loaded. Overall, this hypothetical experiment introduces the concept of both homogeneous and partial stealth multiplets that are susceptible to the sample labelling efficiency, *a*, and the Poisson distribution parameter, *λ*, related to the loading cell rate.

### Defining the four types of multiplets under the multiple sample conditions

We have discussed the prevalence of stealth multiplets in the singleplex condition where cells are classified only as “labelled” or “unlabeled”. However, in the usual mx-scRNA-seq, multiple groups of cells are each stained independently with their own barcodes, and then the multiple groups are pooled together for sequencing. In this configuration, four categories of multiplets can be defined (**Figure 2A**); The first and second categories are the homogeneous and partial stealth multiplets, as seen in the singleplex experiment. The third category is a multilabelled multiplet that consists of cells with more than one barcode, which is detected as a multiplet by sample-barcode demultiplexing and is usually eliminated before further analysis. The last type, an unlabelled multiplet, consists of all unlabelled cells, which is classified as an unlabelled cell-droplet and also easily removed.

**Fig. 2:**
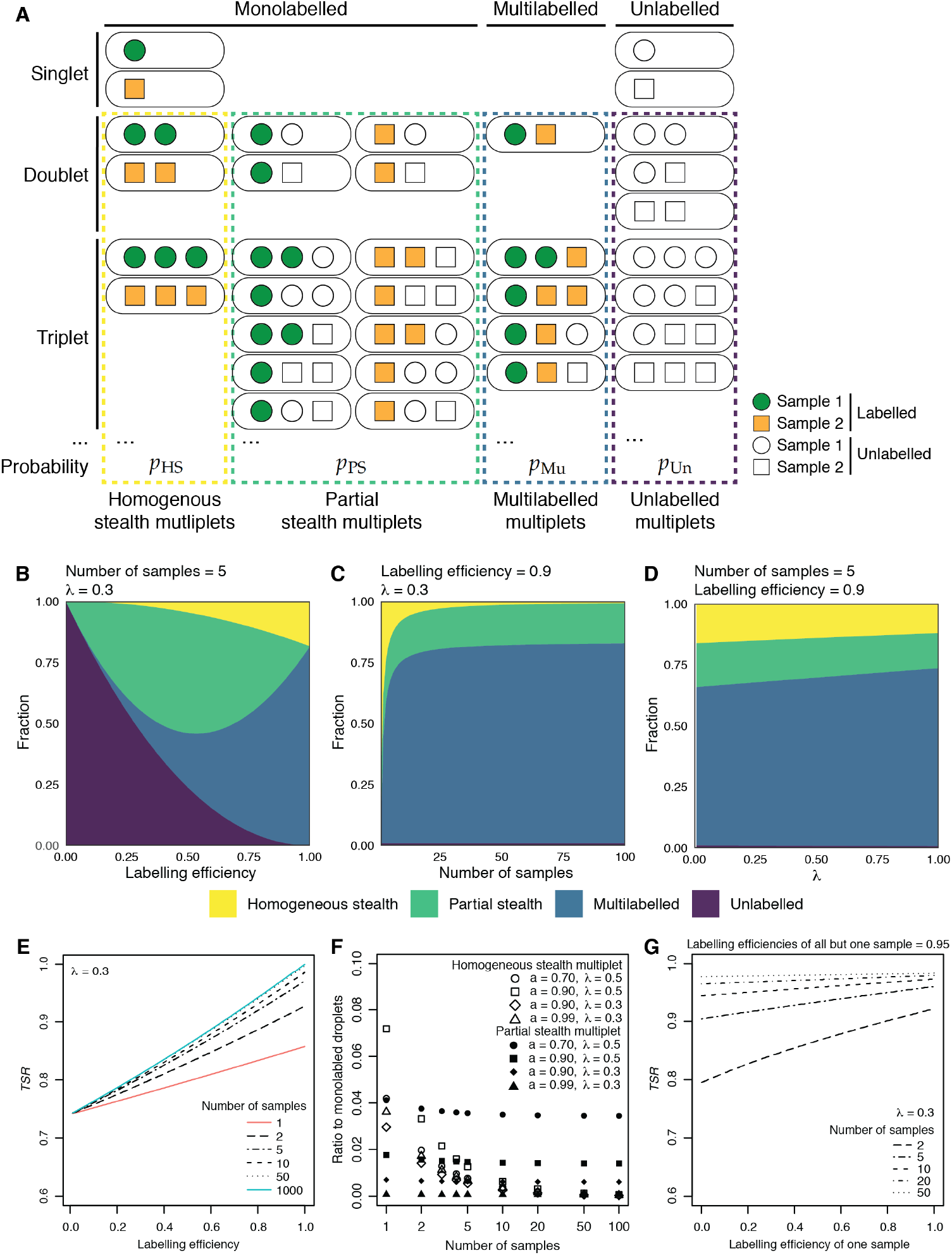
A theoretical probability of the stealth multiplets in a sample-multiplexed experiment. (A) Schematic illustration of the combinations of cells in a single droplet when two samples are multiplexed. Labelled cells are shown in green or orange, depending on the sample labels, and unlabelled in white. The circle and square represent the sample to which cells belong. In the sample multiplexing configuration, the multiplets are classified into four categories: Homogeneous stealth, partial stealth, multilabelled, and unlabelled multiplets, and the parts of the probability for these categories are expressed as *p*_HS_, *p*_PS_, *p*_Mu_, and *p*_Un_, respectively. (B-D) Area charts exhibiting the fraction of the four categories among multiplets when *λ* = 0.3 and five samples are multiplexed (B), when a labelling efficiency is 0.9 and *λ* = 0.3 (C), and when a labelling efficiency is 0.9 and 5 samples are multiplexed (D). (E) *TSR* is plotted against labelling efficiency under various sample numbers. *λ* is fixed at 0.3 (F) The ratio of homogeneous stealth and partial stealth multiplets in the mono-labelled cell-droplets. Note that the partial stealth multiplet persists even when 100 samples are multiplexed. (G) Effects of the labelling efficiency of one sample on *TSR* when *λ* = 0.3 and the labelling efficiencies of the rest of samples are 0.95. All samples are pooled with the same proportion.

The probabilities of each category, *p*_HS_, *p*_PS_, *p*_Mu_, and *p*_Un_ for the homogeneous stealth, partial stealth, multilabelled, and unlabelled multiplet, respectively, are (See **Methods** for the details)

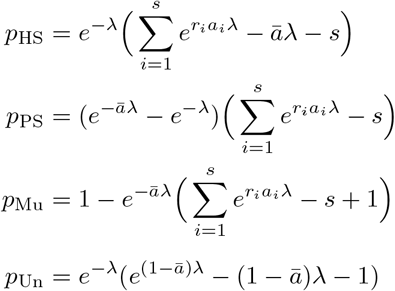

where *s* is the number of samples, *a*_*i*_ is the labelling efficiency of sample *i* (1 *≤ i ≤ s*), comprising a fraction 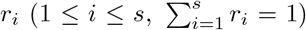 of the pooled population. *ā* is the overall average of labelling efficiencies 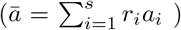, and *λ* is the expected value of the Poisson distribution. Note that *p*_Un_ depends on *ā* because unlabelled multiplets are generated across samples, and the origin of cells consisting of unlabelled droplets cannot be distinguished. Also, *p*_PS_ is expressed both by the overall average *ā* and by the individual labelling efficiency, which supports the idea that the partial stealth multiplets of each sample are formed with unlabelled cell(s) from other samples.

To better understand the characteristics of these probabilities, we shall assume that all samples had an equal labelling efficiency and were pooled with an equal proportion 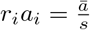 for simplicity. **Figure 2B-2D** show the theoretical fraction of multiplets in eacy category. For a fixed number of samples and a given *λ*, the unlabeled multiplet is the majority when the labelling efficiency is low. As the labelling efficiency increases, the partial stealth multiplet is greater than the multilabelled multiplet for low labelling efficiencies, and then, the homogeneous stealth and multilabelled types become the majority. Ideally, when the labelling efficiency is 100%, the multiplets consist of only the homogeneous stealth and multilabelled types (**Figure 2B**). Interestingly, for a given labelling efficiency and *λ*, the homogeneous stealth multiplets were dramatically reduced as the number of samples increased, but the partial stealth type persists (**Figure 2C**). The value of *λ* does not affect very strongly the composition of the types of multiplets, though it affects the overall multiplet ratio (**Figure 2D**).

We can now define the *TSR* for the multiplex experiment. When *s* samples are multiplexed, the overall *TSR* is given by

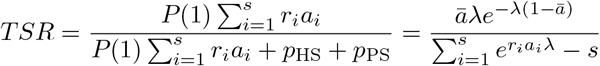

Under the condition 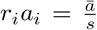 for every *i, TSR* improves as the number of samples increases because a proportion of multiplets can now be identified as multilabelled multiplets (**Figure 2E**). However, the effect of number of samples diminishes when the labelling efficiency is low (**Figure 2E**), because the partial type is the major stealth multiplet. Unlike the homogeneous stealth multiplet, the partial type is not mitigated even when 100 samples are multiplexed (**Figure 2C, 2F**). Lastly, we examine the effect of a poorly labelled sample in pooled population. The poorly labelled sample reduces the overall *TSR* irrespective of the number of multiplexed samples, but this effect can be diminished by increasing the number of samples. This finding suggests that particularly when a few samples are multiplexed, a poorly labelled one may compromise the integrity of the rest of the samples, even though the sample in question can be excluded from the downstream analysis.

### Partial stealth multiplets appeared in a poorly labelled mx-scRNA-seq dataset irrespective of sample demultiplexing methods

So far, we have estimated the theoretical presence and prevalence of the homogeneous and partial stealth multiplet based on the Poisson distribution. Next, we examine an actual mx-scRNA-seq dataset to see if any cells are suspected as partial stealth multiplets, whilst we cannot find potential homogeneous ones without a multiplet remover *in silico*.

We first chose a poorly labelled sample in order to explore our predictions in a case with a high probability of multiplets. The dataset consists of just two samples, from very different cell types so that transcriptome analysis alone will easily distinguish them (providing “ground truth” data): NIH3T3 (3T3) and mouse embryonic stem (ES) cells. Each cell lineage was labelled with a distinct lipid-conjugated oligonucleotide barcode before pooling. Therefore, each population can be identified either by sample barcodes or transcriptome. A dimension reduction plot of the transcriptome data is shown in **Figure 3A**. The three distinct clusters, ES, 3T3, and multiplet of the two lineages, were identified.

**Fig. 3:**
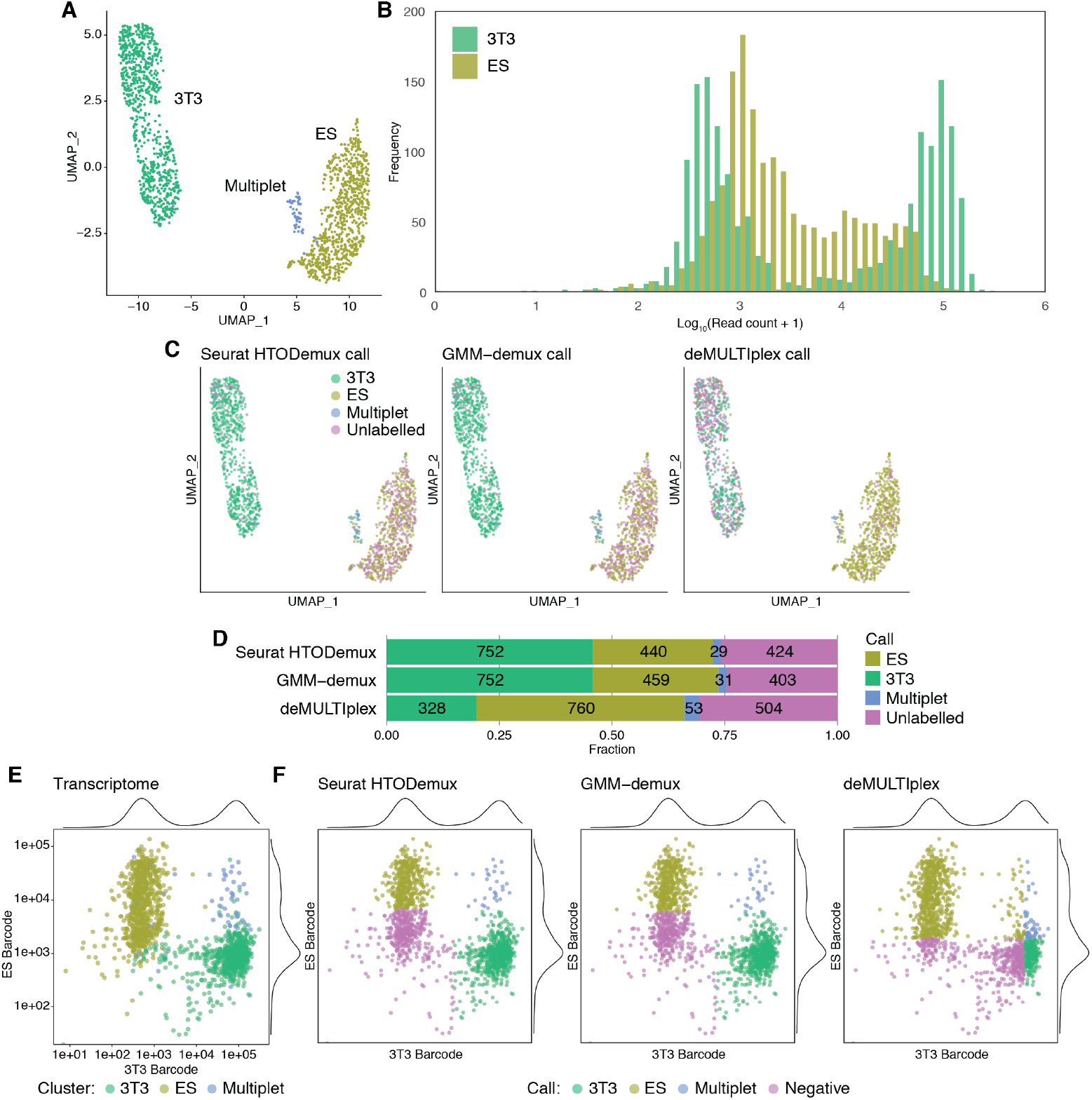
The comparison between cell-type classification by transcriptome and by barcode read demultiplexing in a suboptimally labelled dataset. (A) A dimension reduction plot of the transcriptome data by UMAP. In total, 1,645 cells (803, 784, and 58 cells in the 3T3, ES, and multiplet clusters, respectively) were detected. (B) A histogram of log-transformed barcode read count per cell grouped by each sample barcode. (C) The classification results by the three different algorithms projected on the UMAP of the transcriptome. (D) Summary of the classification results by the three different methods. The cell counts of each group are also shown. (E) A log-transformed scatter plot of the barcode reads colour-coded by the cluster based on the transcriptome. Each barcode density plot (similar to the histogram in (B)) is also shown along each axis. (F) The classification results of each method are projected on the same scatter plots as (E).

A histogram of barcode read per cell illustrated a distinct bimodal distribution for the 3T3 barcode reads, but not for the ES barcode. A low signal-to-noise ratio in the ES barcode reads meant that separating them into two distinct populations was challenging (**Figure 3B**). Next, we applied three different methods of demultiplexing: The HTODemux function from R package Seurat (Seurat::HTODemux) [24], GMM-demux [22], and R package deMULTIplex [14]. **Figure 3C** shows that none of the algorithms could classify the cells based on the sample barcode information as effectively as the “ground truth” transcriptome data. Many of the ES cells were classified as unlabelled cells with Seurat::HTODemux and GMM-demux, and many of the 3T3 cells as unlabelled cells with deMULTIplex (**Figure 3C and 3D**).

Indeed, a scatter plot of the barcode reads illustrated a clear overlap between 3T3 and ES cells in the population around the negative peak of the ES cell barcode reads (**Figure 3E**). Projecting the demultiplexing results on the same scatter plot revealed that Seurat::HTODemux and GMM-demux adopted a conservative threshold in the ES cell classification and lost a substantial part of ES cells and misclassified some of the multiplets as 3T3 cells when compared with the transcriptome cluster (**Figure 3F**). A similar misclassification occurred on the 3T3 cell barcode side by deMULTIplex (**Figure 3F**).

Next, we explored if the misclassified monolabelled cells in the multiplet cluster were partial stealth multiplets. From the clustering by transcriptome, a set of marker genes was defined for each of the ES and 3T3 populations, and a score for each lineage was calculated in each cell. As expected, each population had a high score for its own lineage marker set and showed a good contrast with the other population. Also, the multiplet cluster highly expressed both markers (**Figure 4A**). Interestingly, among the transcriptomic multiplets defined as the double-positive populations for both lineage marker scores, more than half were classified as singlets from just one of the lineages in all demultiplexing algorithms (**Figure 4B**). They were classified as monolabelled cells but had unlabelled cells from the other sample, indicating that they were partial stealth multiplets.

**Fig. 4:**
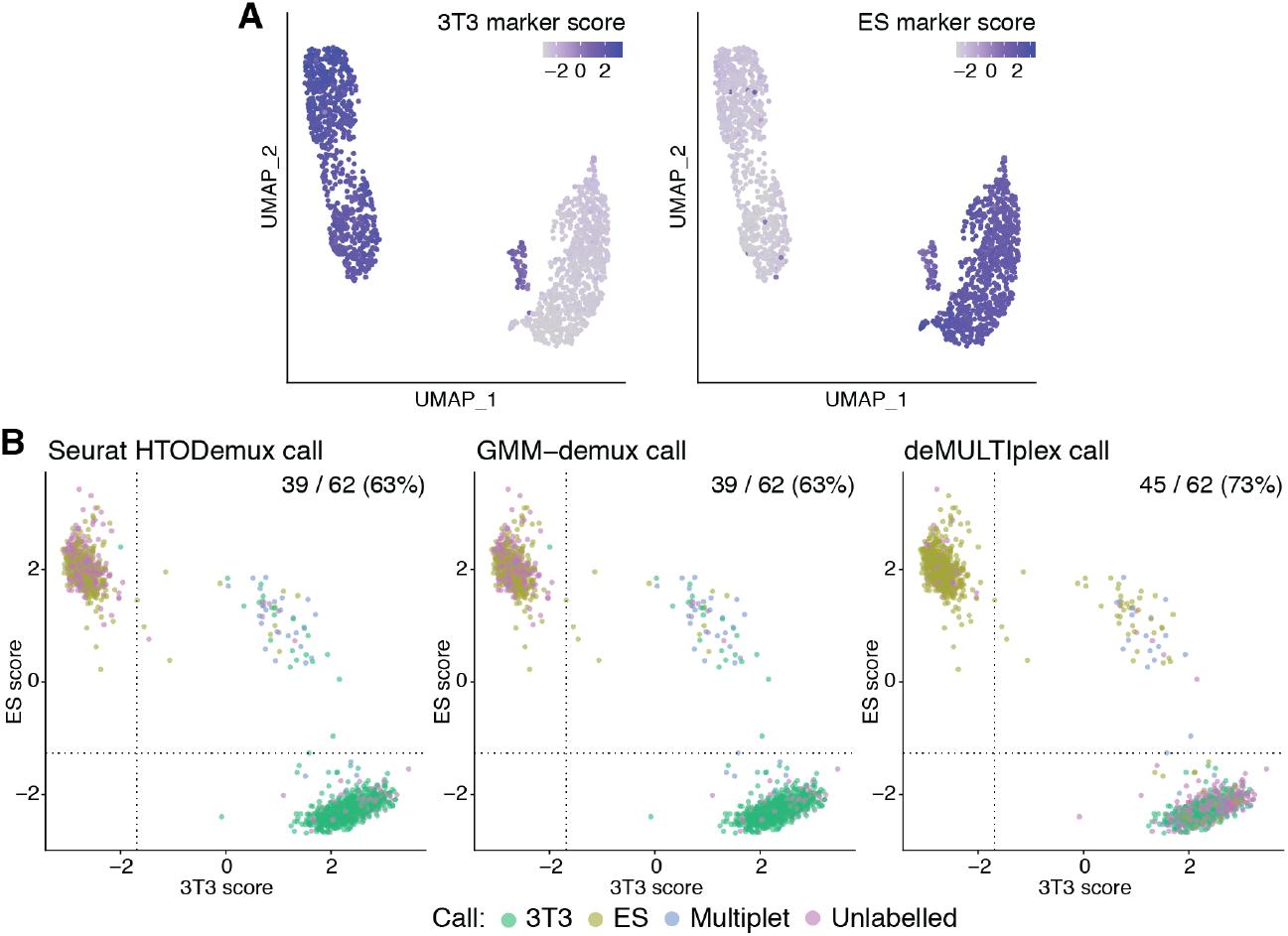
Partial stealth multiplets appear in a suboptimally labelled dataset. (A) The levels of the 3T3 (left) and ES (right) cell marker scores. (B) Scatter plots of the 3T3 and ES scores. The cells in the right upper quadrant were regarded as the multiplets. The number of cells classified as singlets among the multiplets (i.e. partial stealth multiplets), the total number of multiplets, and their percentage are shown at the top right.

To compare the actual results and the theory, we inferred the key parameters, *λ* and the overall average of labelling efficiency *ā* from the demultiplexing results (see **Methods** for details), and estimated the probabilities of the multiplets of each category and *TSR* with **Equations 9 and 10** (2). Notably, the demultiplexing method itself determines each cell’s labelling status, and the apparent labelling efficiency could be different even when analysing the same data set. Also, because the fixed value of *λ* estimated from the specification of the Chromium Controller (See **Supplementary information**) was too small to yield the actual number of the multiplets that appeared in the dataset (We targeted 4,000 cells, it would generate 3.2% of multiplets in theory), we decided to use the inferred value of *λ* as well as the labelling efficiency from the observed cell counts. For reference, multilabelled multiplets, identified as multilabelled cell-droplets, and partial stealth multiplets, identified in **Figure 4B**), from the data are shown in **Table 1**. In this poorly labelled dataset, 2.5% of cell-droplets were expected to be the partial stealth multiplet, which was estimated to be the most prevalent type among the four categories. Indeed, we found at least 2% of cell-droplets were partial stealth multiplets from the data, irrespective of the demultiplexing algorithms (**Table 1**). The partial stealth multiplets in the actual data were underdetected because we could not distinguish the ones consisting of labelled and unlabelled cells from the same sample.

**Table 1:**
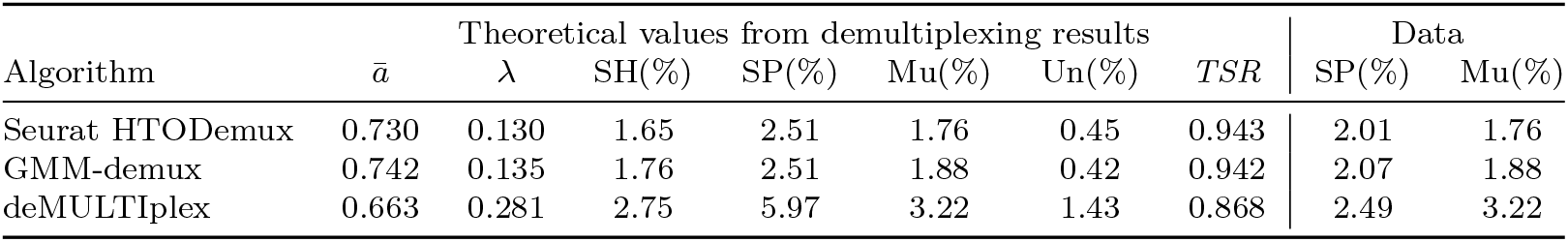
The average labelling efficiency (*ā*), *λ*, and the percentages of each type of multiplets among the whole cell-droplets were estimated from the demultiplexing data shown in **Figure 3**. the observed proportion of the partial stealth multiplet (SP), and the multilabelled multiplet (Mu) are shown on the right. Note that the values of Mu are very close between the theoretical prediction and the observation because we estimated *ā* and *λ* from the observed count of monolabelled, multilabelled, and unlabeled cell-droplets. SH, Homogeneous stealth multiplet. Un, Unlabelled multiplet

From these results, we conclude that when samples are not adequately labelled, it is impossible to identify and exclude the partial stealth multiplet with current demultiplexing methods. Ironically, using a conservative threshold to avoid misclassification increases the risk of the partial stealth multiplet, highlighting the significance of the sample labelling step for a successful mx-scRNA-seq run.

### The partial stealth multiplet was not significant in an optimised experiment and demultiplexing

Lastly, we examined if the partial stealth multiplet is negligible in the properly labelled dataset. The second dataset includes embryonic hind limb bud mesenchyme (HL), ES, and 3T3 cells. Again, each cell lineage was tagged with a distinct sample barcode before pooling. A UMAP plot of the transcriptome data illustrated the three distinct populations (**Figure 5A**). Every sample showed a clear bimodal distribution of barcode read per cell with an altered signal-to-noise ratio, suggesting each cell type has a different affinity with the labelling agent (**Figure 5B**). Interestingly, the demultiplexing results of the three algorithms differed; GMM-demux exhibited the most favourable results, corresponding to transcriptome clusters (**Figure 5C and 5D**). Seurat::HTODemux and deMULTIplex classified many 3T3 cells as unlabelled cells. When the clusters from the transcriptome were projected onto the scatter plot of the barcode reads, the population comprising the high read count peak in each barcode corresponded to each lineage. (**Figure 5E and 5F**). However, the demultiplexing results showed that the Seurat::HTODemux and deMULTIplex set the threshold of the 3T3 barcode high, leading to many negative calls in the 3T3 population (**Figure 5G and 5H**).

**Fig. 5:**
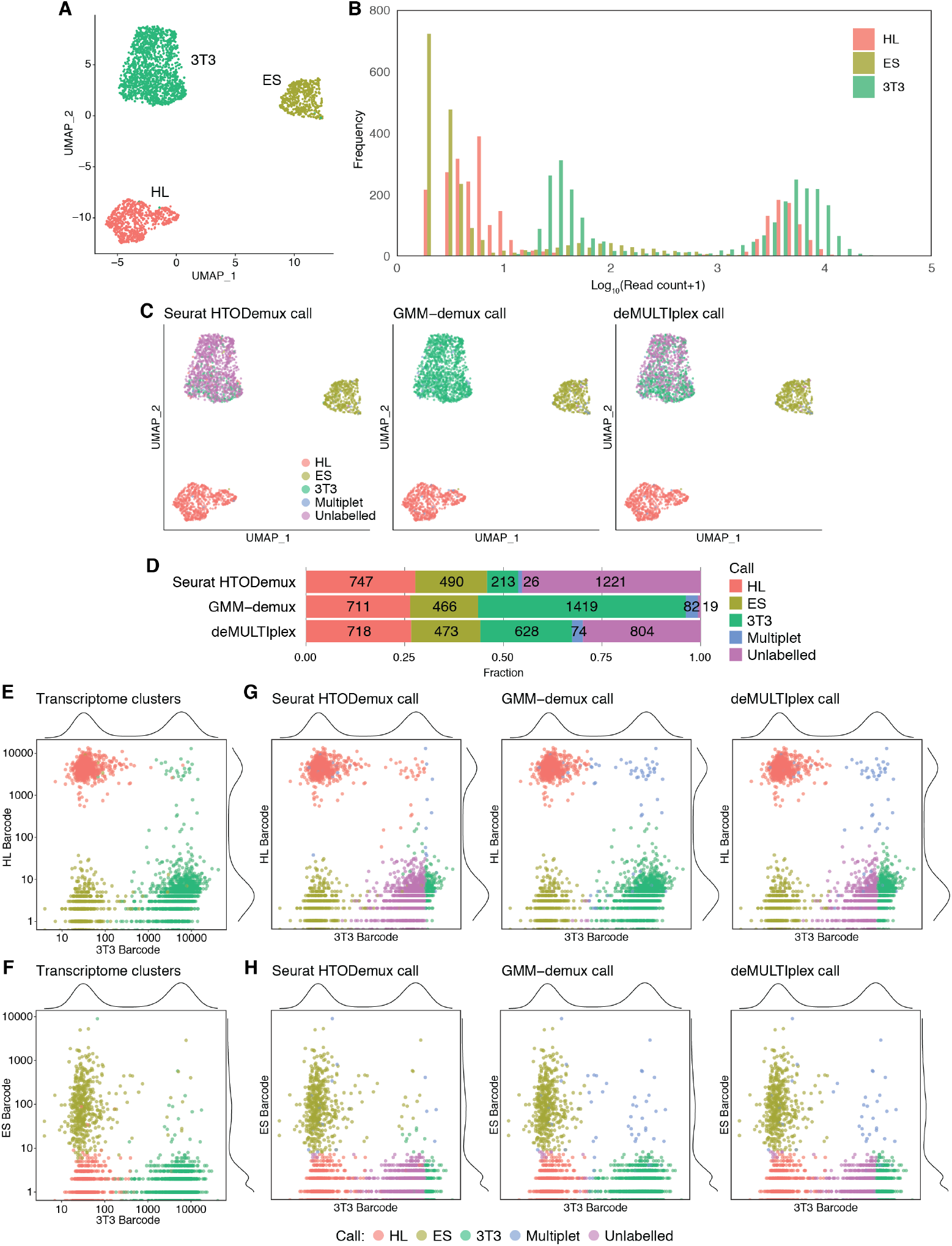
The comparison between cell type classification by transcriptome and barcode read demultiplexing in an optimally labelled dataset. (A) A dimension reduction plot of the transcriptome data by UMAP. In total, 2,697 cells (726, 498, and 1,473 cells in the HL, ES, and 3T3 cell clusters, respectively) were detected. Unlike the two-sample data in **Figure 3**, multiplets belonged to one of the three cell lineage clusters. (B) A histogram of barcode read count per cell grouped by each sample barcode. (C) The classification results of each method are projected on the UMAP plot, as shown in (A). (D) Summary of the classification results by the three different methods. The cell counts of each group are also shown. (E, F) A log-transformed scatter plot of the 3T3 and HL barcode reads (E) and the 3T3 and ES barcode reads (F), colour-coded by the cluster based on the transcriptome. Each barcode density plot (similar to the histogram in (B)) is also shown along each axis. (G, H) The classification results of each method are projected on the same scatter plot as (E) and (F).

The lineage marker scores defined within this dataset showed that most of the cells in each cluster marked high scores on their own lineage marker and low scores on the others, as expected (**Figure 6A**). Notably, several cells showed high scores in other lineage clusters, suggesting they were multiplets with the cells of different lineages (**Figure 6A**). When we examined multiplets based on transcriptome by extracting the cells with positive marker scores in more than one lineage, Seurat::HTODemux yielded a number of the partial stealth multiplet due to the conservative threshold for the 3T3 barcode reads (**Figure 6B-6D**). In contrast, there were few partial stealth multiplets in GMM-demux calls (**Figure 6B-6D**), indicating that the demultiplexing algorithm is another essential factor, and the risk of partial stealth multiplet can be reduced by choosing an adequate method for a given dataset.

**Fig. 6:**
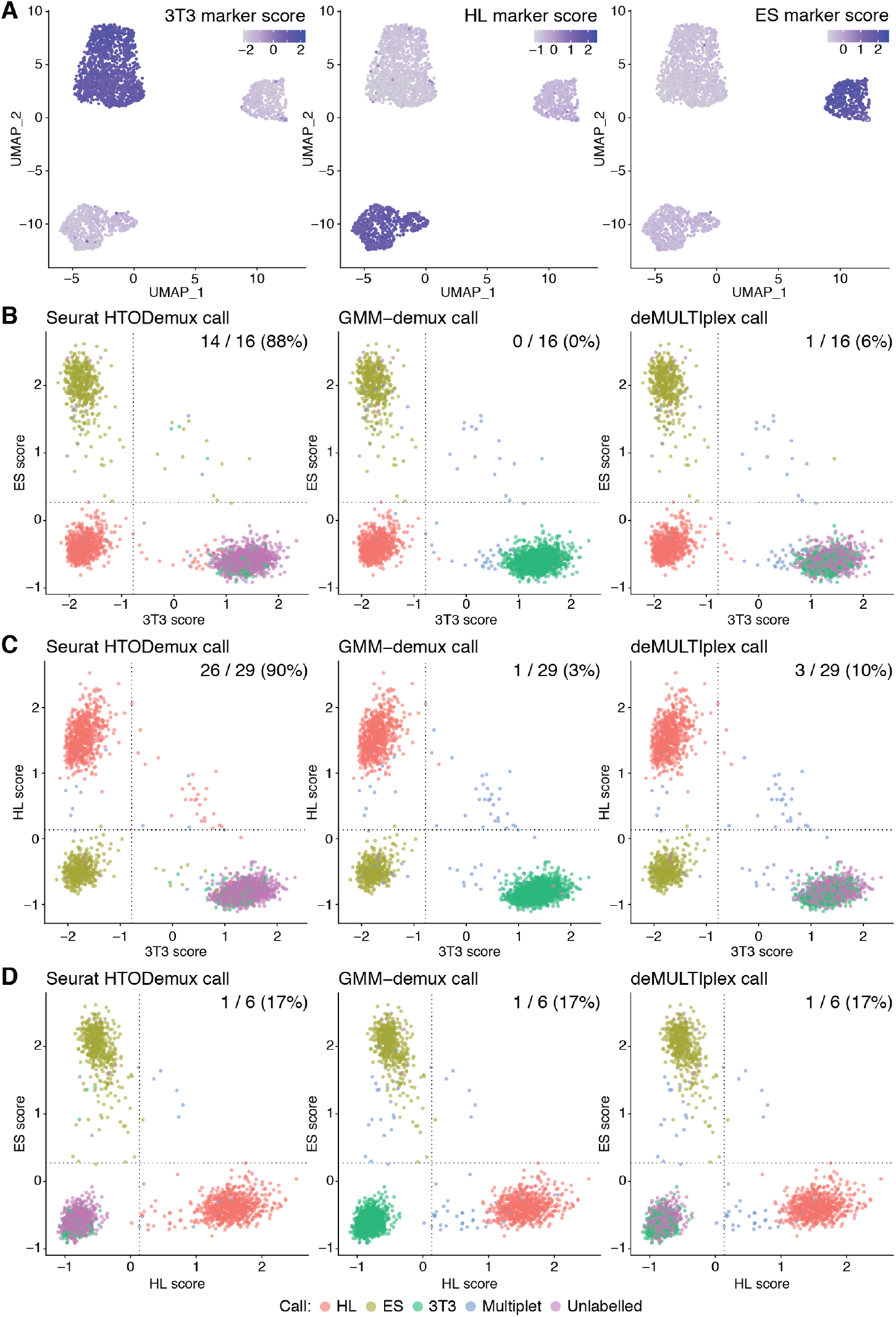
The partial stealth multiplet did not appear in an optimally labelled dataset with an appropriate demultiplexing method. (A) The levels of the 3T3, HL, and ES cell lineage marker scores are shown on the UMAP plot in Figure 5A. (B-D) Scatter plots of the 3T3 and ES scores (B), the 3T3 and HL scores (C), and the HL and ES scores (D). The cells in the right upper quadrant were regarded as the multiplets. The number of cells classified as singlets among the multiplets (i.e. partial stealth multiplets), the total number of multiplets, and their percentage are shown at the top right.

The estimated percentage of multiplets in each category against the total celldroplets is shown in **Table 2**. In this well-labelled dataset (**Figure 5**), the partial stealth multiplet is theoretically only 0.07% with adequate demultiplexing, in this case, GMM-demux. Thus, we concluded the partial stealth multiplet should not be a significant issue in these optimal settings. It is nevertheless striking how different some of the estimated values are from the three different algorithms on the same dataset.

**Table 2:**
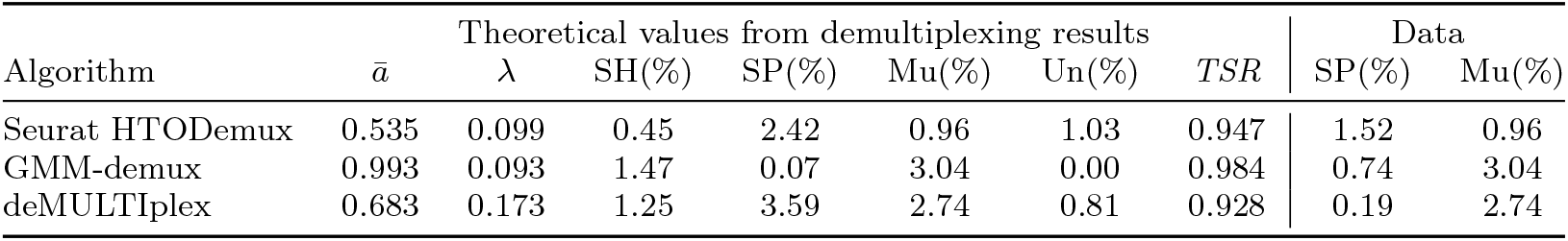
The average labelling efficiency (*ā*), *λ*, and the percentages of each type of multiplet were inferred from the demultiplexing data shown in **Figure 5**, See also the legend of **Table 1**

## Discussion

### The probabilities of homogeneous and partial stealth multiplets and their behaviour

This article provides the first theoretical analysis, as far as we are aware, of the four types of multiplets that occur in mx-scRNA-seq: Homogeneous stealth, partial stealth, multilabelled, and unlabelled multiplets. Among them, the multilabelled and unlabelled multiplets can be excluded as multilabelled and unlabelled cell-droplets. In contrast, the homogeneous stealth and partial stealth multiplets cannot be unambiguously detected because they appear as monolabelled cell-droplets. Therefore, the only solution is high-quality control of the mx-scRNA-seq, which means appropriate sample labelling with a sufficient signal-to-noise ratio *in vitro* and adequate sample demultiplexing *in silico*.

Whilst the homogeneous stealth multiplet, consisting of cells with the same sample barcode, has been pointed out in the previous reports [12, 14, 21, 22], the partial stealth multiplet, harbouring a labelled cell(s) of a given barcode with an unlabelled cell(s), has been overlooked, probably because of the low risk of “label-negative” cells. We examined their probability based on the Poisson distribution model. As a result, both types of multiples depend on the labelling efficiency and *λ*, the mean occurrence of cell-droplet formation event per droplet. Interestingly, although the risk of homogeneous stealth multiplets can be reduced by increasing the number of samples multiplexed, the same approach is not applicable to the partial stealth multiplet because it can be formed with unlabelled cells from any other sample(s).

Mainly two factors determine the predicted labelling efficiency; One is the sample labelling process, and the other is demultiplexing *in silico*. Certainly, optimising the labelling process to maximise the signal-to-noise ratio is essential. On the other hand, whether these labelled cells are classified as label-positive depends on how the algorithm differentiates the positive and negative populations. In other words, the demultiplexing algorithm *per se* defines the apparent labelling efficiency by cutting off the continuous barcode read counts. Therefore, choosing an appropriate demultiplexing method is another critical step in mx-scRNA-seq.

The other key parameter, *λ*, which is correlated with cell loading rates onto a microfluidic device, determines the overall multiplets ratio. Therefore, loading fewer cells can reduce the risk of overall multiplets. However, the recent trend in mx-scRNA-seq is increasing the number of loading cells by expecting effective multiplet removal [14, 21]. Although increasing the number of sample barcodes can effectively remove homogeneous stealth multiplets and multilabelled multiplets, it cannot reduce partial stealth multiplets. Therefore, the prediction here will be a caveat against the suboptimally labelled mx-scRNA-seq, especially when the excessive number of the cells are “super-loaded.” Additional use of the multiplet removal program *in silico* is suggested, particularly when a small number of samples are multiplexed (against the homogeneous stealth multiplets), when low labelling efficiency is predicted (against the partial stealth multiplets), or when the *λ* is expected to be high (against both types of stealth multiplets), if those programs’ assumptions hold true in the data in question.

### Choosing an appropriate demultiplexing method

As shown in **Table 2**, the demultiplexing algorithms’ calls were different even in the optimally labelled dataset.; there was a significant difference in the classification of 3T3 cells, highlighting the importance of choosing an appropriate demultiplexing method. We observed that the conservative threshold of parameters ironically increased the prediction of partial stealth multiplets by yielding more unlabelled cells. Therefore, It is important to demultiplex the cells with an algorithm based on a relevant sample-barcode read distribution model.

Many sample-demultiplexing methods have been proposed so far: CITE-seq [13], Seurat::HTODemux [24], deMULTIplex [14], GMM-demux [22], BFF [25], demuxmix [26], and deMULTIplex2 [27]. In our case, GMM-demux was optimal for the well-labelled dataset, which is probably because the dataset has only three samples with a sufficient read depth to capture the bimodal log-standard distributions. On the contrary, deMULTIplex2 [27], which is one of the latest algorithms, is not applicable because our datasets violate their assumption in terms of the cell size variation among samples (Qin Zhu, personal communication), which emphasises the importance of exploring an appropriate method utilising a relevant assumption for a given dataset.

### Limitation of the Poisson distribution modelling and further consideration

In this article, we modelled the probabilities of multiplets in scRNA-seq based on the Poisson distribution of which assumptions apparently fit well, and indeed multiple previous reports utilised it to discuss the probability of cell-droplet formation [6, 10, 22, 23]. Still, there are limitations to using this model. The first point is that estimating the key parameter *λ* in routine experimental settings is challenging because we cannot see the total number of droplets. The *λ* from the specification of the microfluidic device may be inaccurate due to a technical variation of devices and experiments, particularly cell counting steps. In this study, the *λ* estimated from the specification of microfluidic device was apparently small to yield the resulted number of multiplets, and we inferred *λ* from demultiplexing results.

The Poisson distribution assumes randomness, which means the previous event did not affect the subsequent cell-droplet formation. However, there may be some minor deviations from this assumption; The multiplet cells that contacted each other due to incomplete dissociation violate this assumption and distort the multiplet ratio. Also, the Poisson distribution assumes the expected value, *λ*, is constant, and in this paper, we applied this assumption to the cell-droplet formation. However, there is another factor - the beads for capturing transcripts. And we can observe the cells only when a bead exists in the same droplet. In the case of the Chromium Controller (10X Genomics), a report showed that 16.1% had no beads, 80.0%, and 3.9% of the droplet had 0, 1, and *≤*2 beads, respectively [28]. Bead integration is much more frequent than that of cells, but it does not always happen, and thus, this may be a source of error in estimating the multiplet ratio. Another report showed that when a very high number of cells were loaded, there was a discrepancy between the Poisson distribution and observed cell counts, and negative binomial distribution better explained the actual cell-droplet formation, suggesting that the expected value *λ* is not constant in the actual cell-droplet formation [29]. Nonetheless, the Poisson distribution model can concisely describe the process of cell-droplet formation and is useful for estimating the probability of each type of multiplets within a range of cell loading rates in most mx-scRNA-seq experiments.

## Conclusion

There are four types of multiplets in mx-scRNA-seq: Homogeneous stealth, partial stealth, multilabelled, and unlabelled multiplets. Among them, the partial stealth multiplet, which consists of a monolabelled cell(s) with an unlabelled cell(s), has not previously been scrutinised. We have theoretically quantified this category and illustrated its presence in real actual datasets. Under optimised experimental settings, the partial stealth multiplets are few. However, when either sample labelling *in vitro* or demultiplexing *in silico* is suboptimal, it may become a major type of multiplets and compromise the integrity of the dataset irrespective of the number of multiplexed samples. These results will be a caveat against an experiment without optimising the sample labelling step. Also, examining multiple demultiplexing algorithms is recommended.

## Methods

### Probabilities of stealth multiplets in the singleplex condition

Under the singleplex experimental condition (**Figure 1A**), the probability of empty droplets, singlets, and multiplets are expressed as *P* (0), *P* (1), and *P* (*k ≥* 2), respectively. In addition, the multiplets can be divided into homogeneous stealth, partial stealth, and unlabelled multiplets (**Figure 1A**). Therefore, the whole events in this configuration are,

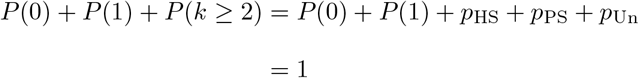

where *p*_HS_, *p*_PS_, and *p*_Un_ are the parts for the probability of homogeneous stealth, partial stealth, and unlabelled multiplets. First, *p*_HS_ can be expressed as

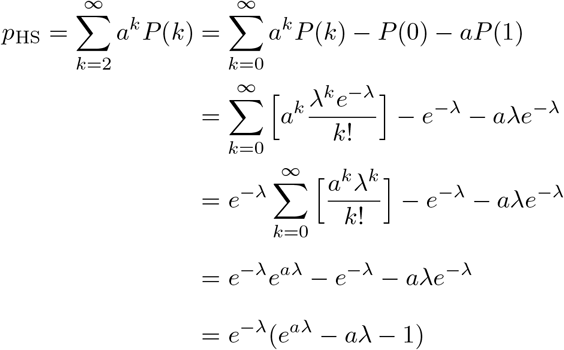

where *P* (*k*) is a Poisson distribution function, *λ* is the average chance of the event per droplet in which a cell enters into a droplet (0 *< λ <* 1), and *a* is a labelling efficiency, which means a fraction of labelled positive cells (0 *≤ a ≤* 1).

Similarly, *p*_Un_ is

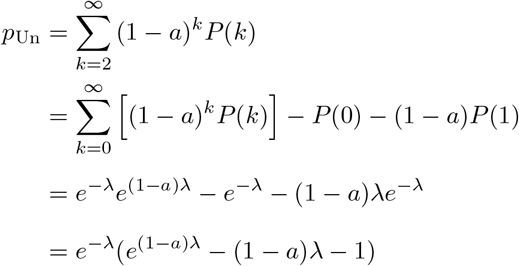

Therefore, *p*_PS_ is

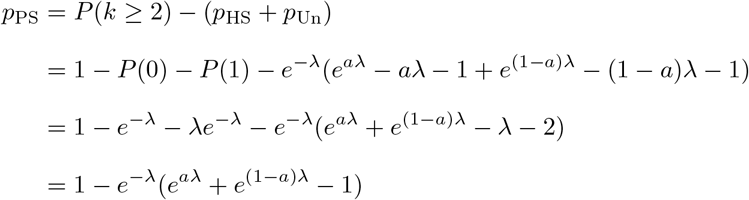

The probability of stealth multiplet is *p*_HS_ + *p*_PS_ and expressed as

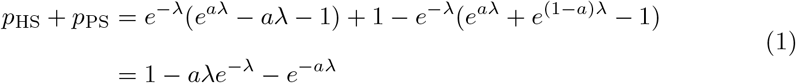

Therefore, the ratio of true singlets among the monolabelled cells (*TSR*) is

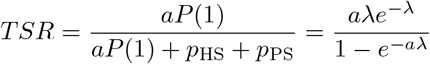

### Probabilities of stealth multiplets under the multiple sample condition

In the singleplex experiment, we did not consider the number of samples. Next, we take the samples with different barcodes each other into account (**Figure 2A**). Here, we examine the probabilities of the four categories of multiplets. In addition to the three categories that appeared in the singleplex case, we introduce *p*_Mu_, the parts of the probability for the multilabelled multiplets that are only generated when multiple sample barcodes are pooled. Therefore, the whole events are

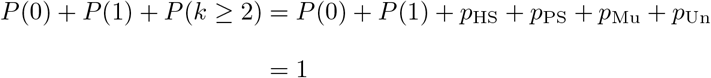

Given *s* samples are labelled with a different barcode per sample under a labelling efficiency, *a*_*i*_ (1 *≤ i ≤ s*, 0 *≤ a*_*i*_ *≤* 1), and *s* samples are pooled with a proportion of 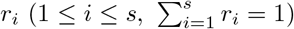, the overall average of labelling efficiencies, *ā*, is

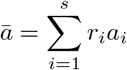

First, *p*_HS_ is expressed as the sum of homogeneous stealth multiplets of each sample as follows:

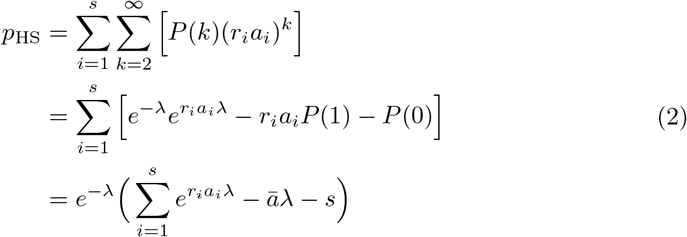

On the other hand, *p*_Un_ can be expressed as

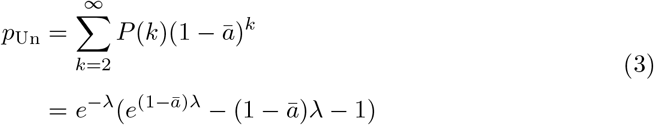

where *ā* is the whole average of the labelling efficiency, because unlabelled multiplets are generated across samples, and we cannot distinguish the origin of cells consisting of unlabelled cell-droplets.

Next. we put the fraction of the homogeneous stealth multiplet, *f*_HS_(*k*), and the unlabelled multiplets, *f*_Un_(*k*), in each *k*-et cell-droplets.

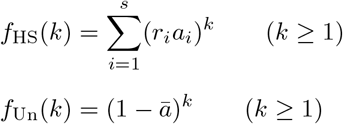

Here, we regard the labelled singlets and unlabelled singlets as special cases of homogeneous stealth multiplets and unlabelled multiplets, respectively, and extend the definition of *f*_HS_(*k*), and *f*_Un_(*k*) to *k* = 1. Note that these true singlets are not included in *p*_HS_ and *p*_Un_. Then, the fraction of the partial stealth multiplet in *k*-et cell droplets, *f*_PS_(*k*), can be described as the sum of the combinations of the fraction of homogeneous stealth *j*-et multiplets, *f*_HS_(*j*), and unlabelled (*k − j*)-et multiplets, *f*_Un_(*k − j*) (1 *≤ j ≤ k −* 1). That is, *f*_PS_(2) is a combination of *f*_HS_(1) and unlabelled singlets, *f*_Un_(1). *f*_PS_(3) is the sum of the combination of *f*_HS_(1) and *f*_Un_(2), and *f*_HS_(2) and *f*_Un_(1). Similarly, *f*_PS_(*k*) is the sum of the combination products of *f*_HS_(1) and *f*_Un_(*k −* 1), *f*_HS_(2) and *f*_Un_(*k −* 2), …, and *f*_HS_(*k −* 1) and *f*_Un_(1), and expressed as

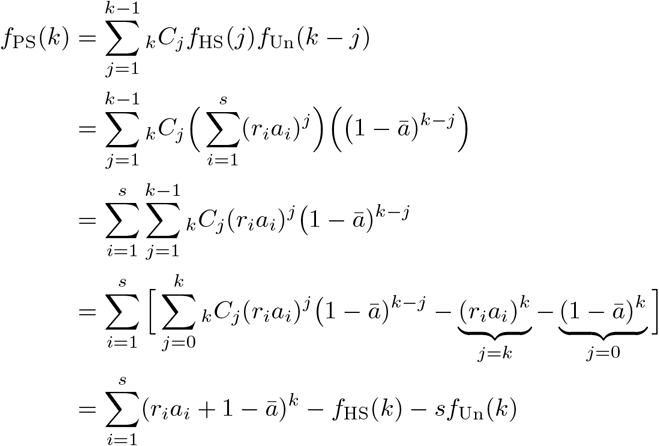

From this equation, the probability of both types of stealth multiplets, *p*_HS_ + *p*_PS_ is expressed as follows:

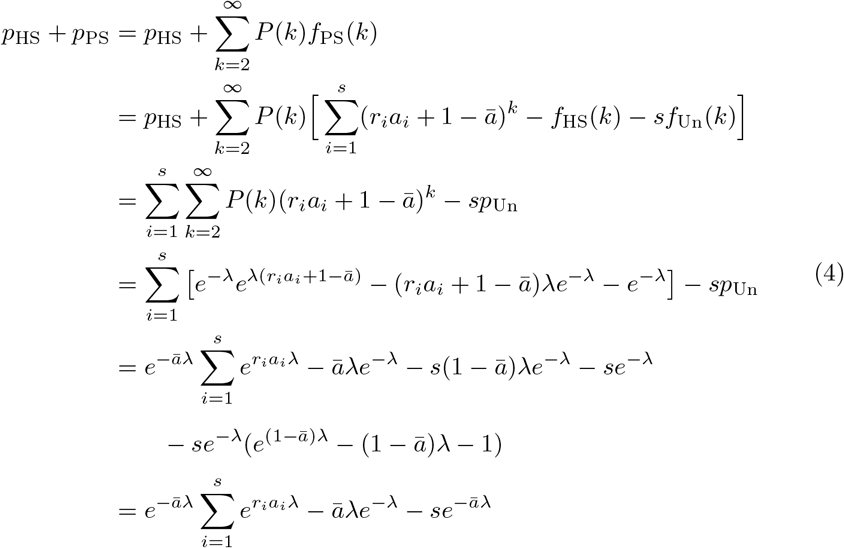

Note that in the singleplex case, which means *s* = 1, *r*_*i*_*a*_*i*_ = *ā*, the probability of the stealth multiplet is

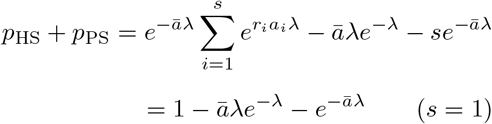

which is equal to **Equation (Eq.) 1**. From the **Eq. 2, 3, and 4**, we can express the probability of the other categories of multiplets. First, the probability of partial stealth multiplet *p*_PS_ is

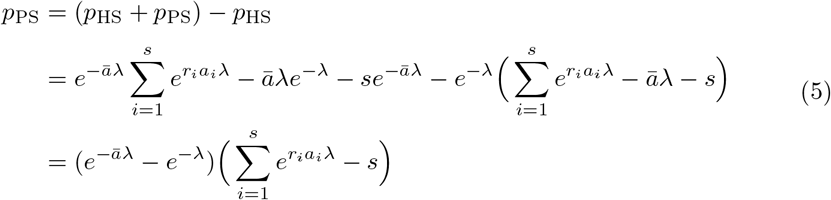

Then, the probability of multilabelled droplet *p*_Mu_ can be expressed as the complement of the other three types of multiplets (*p*_Mu_ = 1 *− P* (0) *− P* (1) *−* (*p*_HS_ + *p*_PS_ + *p*_Un_)). From **Eq. 3 and 4**, the sum of *p*_HS_, *p*_PS_, and *p*_Un_ is expressed as

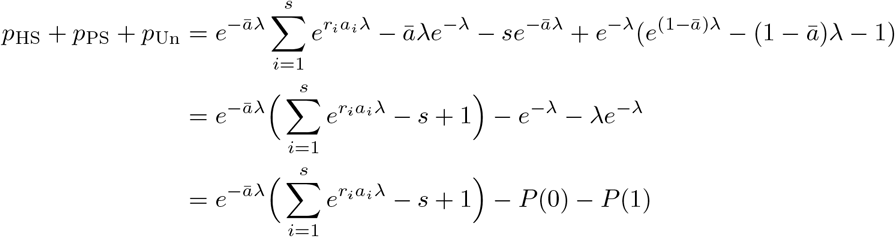

Therefore, the probability of the multilabelled multiplet is

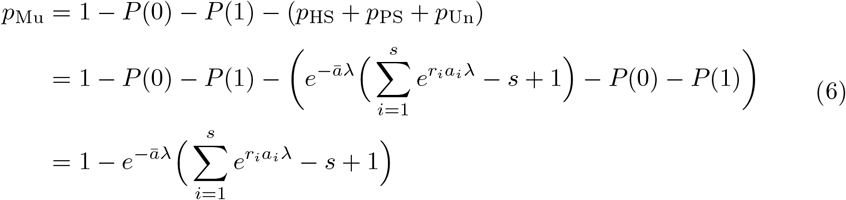

Lastly, when *s* samples are multiplexed, the overall *TSR* is

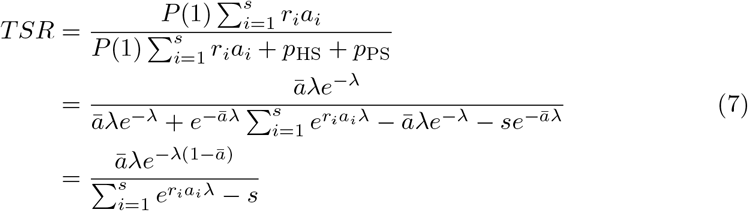

### Assuming the special case in which all samples are labelled with an equal efficiency and pooled with an equal proportion

To facilitate the exploration of the behaviour of the probability of the multiplets in **Figure 2B-2F and Table 1 and 2**, we adopted a special case in which all samples are labelled evenly and pooled with the same proportion. in this case, *a*_*i*_ = *ā* and *r*_*i*_ = 1 */ s* for given *i*, and thus,

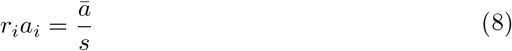

Under this condition, the probability of each category (**Eq. 2, 5, and 6**) can be expressed as,

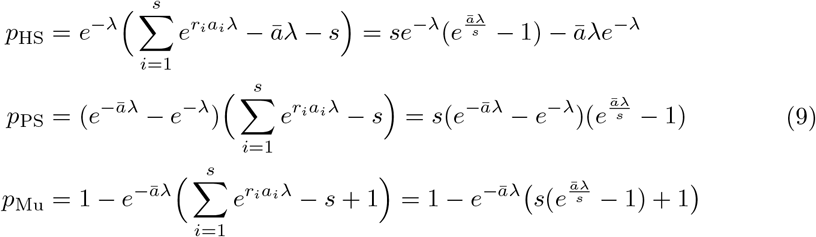

From these equations, together with the probability of the unlabelled multiplet (**Eq. 3**), we can explore their behaviour by considering only *ā* and *λ*.

Lastly, from **Eq. 7**, the *TSR* under this condition is

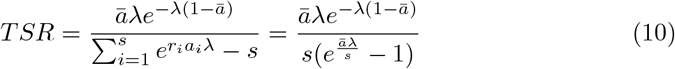

### Estimating *λ* and a labelling efficiency from demultiplexing results

Because the total droplet count is unknown, the only clues to estimate the parameters, *λ* and *ā*, the overall average of labelling efficiencies, are the ratio of observed cell numbers, particularly that of monolabelled/unlabelled cell-droplets. First, from **Eq. 3 and 4**, the probability of monolabelled and unlabelled cell-droplets are

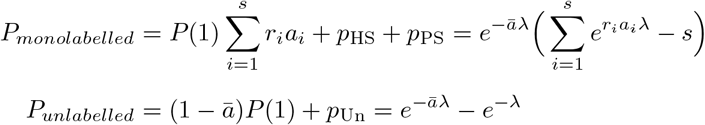

Therefore, the ratio of monolabelled / unlabelled cell-droplets is

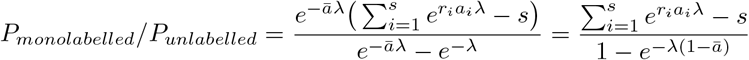

By assuming that the labelling efficiencies of individual samples are equal, and the equal proportion of cells are pooled, **Eq. 8** is applied, and we can approximate the ratio as

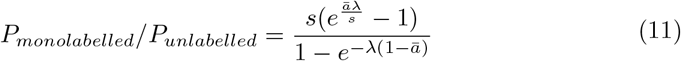

In addition, we can also observe the total number of detected cells, which enables us to introduce another ratio, unlabelled cell-droplets to total ones. Because the probability of the total number of cell-droplets is expressed as 1 *− P* (0) = 1 *− e*^*−λ*^, the ratio is

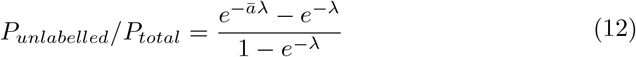

Now we have two equations (**Eq. 11 and 12**) for the two parameters, *λ* and *ā*. In this article, we numerically solved for *λ* and *ā* by the Newton-Raphson method with the multiroot function of rootSolve package (version 1.8.2.3) in R (version 4.1.2). The initial condition for *ā* was given by

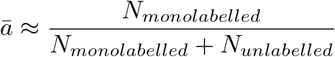

where *N*_*monolabelled*_ and *N*_*unlabelled*_ were the numbers of monolabelled and unlabelled cells from the demultiplexing results, respectively. This value corresponds to an estimated *ā* based on singlets alone. For *λ*, we used 0.05 as an initial condition and increased it by 0.05 until we obtained a positive root. The estimated results for the dataset in **Figure 3 and 5** are shown in **Table 1 and 2**, respectively.

### Graphical representations

All graphical illustration was done with R (version 4.1.2) [30], and linear regression was performed with the stats::lm function.

### Mice

All mice were bred at the PRBB Animal Facility, which is accredited by the International Association for Assessment and Accreditation of Laboratory Animal Care (AAALAC), and handling and care were done in accordance with the AAALAC criteria. All procedures in this study were approved by EMBL Institutional Animal Care and Use Committee (IACUC), and mouse embryos were collected by following the American Veterinary Medical Association (AVMA) guidelines. The B6.129-*Sox9*^*tm1Haak*^ (Sox9-EGFP) mice (MGI:3844977)[31] were maintained by backcrossing to C57BL/6J mice.

### scRNA-seq

#### Sample preparation

A hindlimb bud’s distal region corresponding to the future autopod was obtained from a Sox9-EGFP mouse embryo at E11.5. The cell attachements were loosened by incubating with TrypLE Express (Thermo Fisher Scientific) at 37°C for 15 minutes, and the ectoderm was removed under the stereoscope. The isolated mesenchyme clump was put in a Cell Prep Buffer (Ca^2+^, Mg^2+^-free phosphate-buffered saline (PBS) with 0.1% poly(vinyl alcohol) (PVA) and 1 mM ethylenediaminetetraacetic acid (EDTA)), dissociated by gentle pipetting. NIH3T3 cells were cultured with High glucose Dulbecco’s Modified Eagle Medium (DMEM) (Thermo Fisher Scientific) supplemented with 10% Fetal Bovine Serum (FBS), L-Glutamine, and Penicillin/Streptomycin. The transgenic T-p2a-H2B-eGFP and CAG::H2B-mKO2 Mouse ES cells (unpublished) derived from G4 [32] ES cells were cultured with DMEM supplemented 10% FBS and leukemia inhibitory factor (LIF) as described [33]. Both cultures were dissociated with TrypLE Express at 37°C for 3 minutes, following multiple washes with PBS to remove serum. The reaction was stopped by adding four volumes of a Wash Buffer (1:1 mixture of DMEM and Ham’s F12 medium (Thermo Fisher Scientific) with 0.1% bovine serum albumin (BSA)) to TrypLE Express, and cells were completely dissociated by gentle pipetting. After pelleting the cells by centrifugation, they were resuspended in the designated buffer for sample labelling as described below.

#### MULTI-seq

The three-sample-multiplexed dataset (**Figure 5 and 6**) was generated with MULTI-seq[14]. The hindlimb bud (HL) cells, ES cells, and NIH3T3 cells were resuspended in a Cell Prep Buffer and filtered through a 35-µm cell strainer cap. After counting cells, 1*×* 10^5^ cells were pelleted and resuspended in 180 µL of Cell Prep Buffer. These cell samples were labelled with MULTI-seq barcode oligonucleotides as described [14]. Briefly, a 1:1 mixture of the cholesterol-conjugated Anchor-oligonucleotides (Anchor CMO, synthesised by Integrated DNA Technologies) and Barcode oligonucleotides with a distinct barcode sequence for each sample (final concentration, 0.2 µM) was added to each cell sample and incubated on ice for 5 minutes. Next, the same concentration of Co-Anchor CMO (synthesised by Integrated DNA Technologies) was added and incubated for another 5 minutes. After labelling, the labelled cells were washed with PBS with 1% BSA (PBS-BSA (1%)) three times, resuspended in 100 µL of PBS-BSA (1%), and pooled. The pooled sample was filtered through a 35-µm cell strainer and counted again before loading onto the microfluidic chip described below. The sample-barcode sequences for the HL, ES, and NIH3T3 cells were “CATAGAGC”, “TCCTCGAA”, and “GTGTACCT”, respectively.

The pooled single-cell suspension was encapsulated with Chromium Single Cell 3’ GEM, Library & Gel Bead Kit v2 (10X Genomics) and a Chromium Single Cell Chip A Kit (10X Genomics) by the Chromium Controller (10X Genomics, firmware version 4.00) and transcripts and MULTI-seq barcode oligos were captured and reverse-transcribed as described in the manufacturer’s instructions. We targetted 4000 cells to sequence. The 3’-enriched cDNA library construction were performed with Chromium Single Cell 3’ GEM, Library & Gel Bead Kit v2 (10X Genomics) according to the manufacturer’s instructions. The MULTI-seq barcode oligos were fractionated after the cDNA amplification, amplified and indexed by PCR with KAPA HiFi HotStart ReadyMix (Roche) for sequencing. Both cDNA and MULTI-seq-barcode libraries were sequenced with NextSeq500 (Illumina). We read 8 base pairs (bp) for TruSeq Indices, 26 bp for 10X barcodes and unique molecular identifiers (UMIs), and 58 bp for both fragmented cDNA and MULTI-seq barcodes.

#### 3’ CellPlex

For the two-sample-multiplexed dataset (**Figure 3 and 4**), sample labelling was done with 3’ CellPlex Kit Set A (10X Genomics), according to the manufacturer’s protocol. Briefly, the single-cell suspension of the NIH3T3 cells and ES cells were resuspended in PBS-BSA (0.04%) and filtered through a 35-µm cell strainer. After counting cells, 1.9*×*10^6^ of NIH3T3 cells and 1.6*×*10^5^ of ES cells were collected by centrifugation and resuspended in 100 µL of Cell Multiplexing Oligo (CMO) 301 and CMO 302, respectively. After 5 minutes of incubation on ice, samples were washed with PBS-BSA (1%) three times and resuspended in PBS-BSA (1%), followed by cell counting. These samples were pooled so that the pooled sample contained equal numbers of cells from each cell type. The pooled samples were again filtered with a 35-µm cell strainer before loading onto the microfluidic chip. Cell encapsulation and barcoding were performed with 3’ Next GEM Single Cell 3’ Kit v3.1 (10X Genomics) and a Chromium Single Cell Chip G Kit (10X Genomics) by the Chromium Controller (10X Genomics, firmware version 4.00) according to the manufacturer’s protocol. The 3’-enriched cDNA and Feature Barcodes libraries were prepared with 3’ Next GEM Single Cell 3’ Kit v3.1 (10X Genomics) and Feature Barcode Kit (10X Genomics) following the manufacturer’s instructions. These libraries were sequenced with NextSeq2000 (illumina). We read 10 base pairs (bp) for i7 and i5 indices, 28 bp for 10X barcodes and UMIs, and 90 bp for both fragmented cDNA and Feature Barcodes.

## Data analysis

### Quality control of scRNA-seq

The sequencing reads were aligned and mapped on the reference mouse genome (mm10) and counted for each feature to build a feature-cell matrix with CellRanger count (version 6.0.1, 10X Genomics) for MUTLI-seq or CellRanger multi (version 6.1.1, 10X Genomics) for 3’CellPlex. The filtered feature-cell matrix from the CellRanger was used for MULTI-seq, and the raw feature-cell matrix was used for 3’CellPlex, which was further screened with the R package DropletUtils (version 1.14.2) [34] for cell-droplet candidates. These feature-cell matrices were further screened for quality control with the number of genes detected and mitochondrial gene reads (**Supplementary Figure 1**). Next, the gene expression levels were normalised with the scater (version 1.22.0) [35] and scran (version 1.22.1) [36] packages from R (version 4.1.2) [30].

### Clustering and visualisation

The normalised data were further analysed to remove a technical variance by the Poisson noise model, and biological variance for each gene was estimated with the scran package (version 1.22.1) [36]. Highly variable genes (HVGs) for dimension reduction were defined as the genes with a biological variance *>*0.1 (1,714 genes) and *>*0.3 (2,461 genes) for the three-sample and the two-sample dataset, respectively. After data standardisation with the ScaleData function in the Seurat package (version 4.3.0) [24], principal component analysis (PCA) was done on the selected HVGs, followed by Uniform Manifold Approximation and Projection (UMAP) by using the first 20 dimensions from the PCA with the Seurat package in R. Clustering was done with the FindCluster function in the Seurat package with the default original Louvain algorithm with the resolution 0.1 and 0.01 for the three-sample and the two-sample dataset, respectively.

**Supplementary Figure 1:**
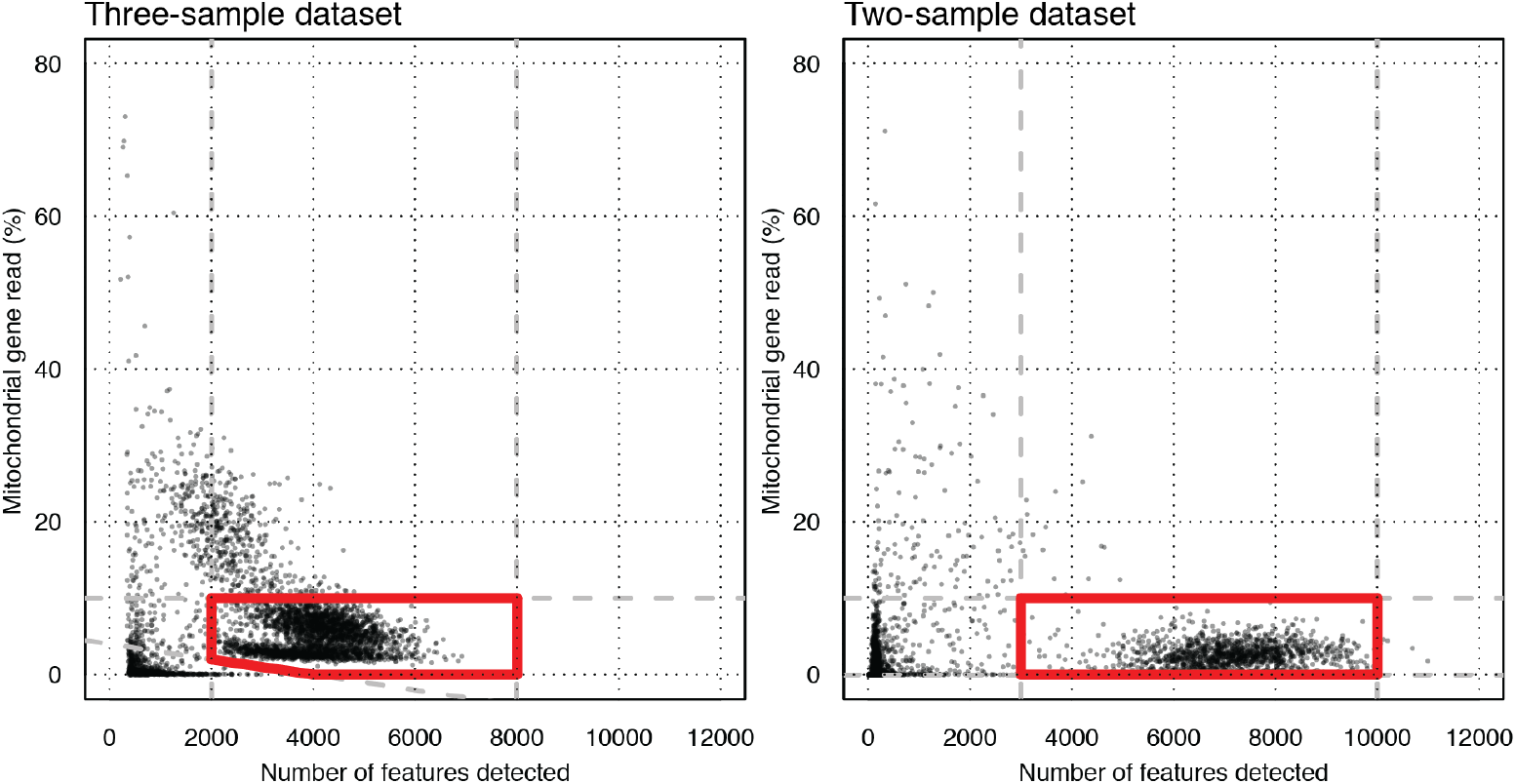
The QC filter for scRNA-seq. Scatter plots showed the number of features detected and the ratio of mitochondrial gene reads (%). The cells plotted inside the red polygons were filtered for further analysis.

### Sample Barcode demultiplexing

For MULTI-seq, counting sample barcodes and building a cell-barcode read count matrix were done with the MULTIseq.preProcess and MULTIseq.align functions of the deMULTIplex package (version 1.0.2) in R (version 4.1.2). For the 3’CellPlex, the Feature Barcode tag was counted with CellRanger multi (version 6.1.1, 10X Genomics). We examined three different demultiplexing algorithms:

deMULTIplex (version 1.0.2) [14] in R (version 4.1.2): After one round of the findThresh and classifyCells functions, removing the label-negative cells and repeated it as the second round—the classification after two rounds were used as final calls.

Seurat::HTODemux (version 4.3.0) [24] in R (version 4.1.2): First, each barcode count was normalised with centred log-ratio (CLR) transformation with the NormalizeData(normalization.method = “CLR”) function. Then cells were classified with the HTODemux(kfunc = “clara”, positive.quantile = 0.99) function based on the normalised barcode matrix.

GMM-demux (version 0.2.1.3) [22]: Demultiplexing was done with the default confidential score threshold, -t 0.8, and designating expected number of cells by -u 2697 and -u 1645 options for the three-sample and the two-sample dataset, respectively.

### Detecting the partial stealth multiplet

The marker genes for each transcriptome cluster were defined with the FindMarkers function in the Seurat package (version 4.3.0). We adopted the top 25 differentially expressed genes as marker genes and calculated each marker score with the AddModuleScore function in the Seurat package (version 4.3.0) function. We fitted a two Gaussian curves to the score and set the threshold as *p* = 0.0001 estimated from the average of the lower curve with the twoGaussiansNull(p.adj.method = “BY”, max.adj.p = 0.0001) function of the Ringo package (version 1.58.0) [37] in R. The cell-droplet with the scores above the thresholds for more than one lineage was defined as a multiplet from the transcriptome. Among them, the ones classified as monolabelled cell-droplets were judged as partial stealth multiplets.

## Supplementary information

Supplementary material file (SupplementaryMaterial.pdf) is attached with this paper.

## Ethics approval and consent to participate

All mice were bred at the PRBB Animal Facility, which is accredited by the International Association for Assessment and Accreditation of Laboratory Animal Care (AAALAC), and handling and care were done in accordance with the AAALAC criteria. All procedures in this study were approved by EMBL Institutional Animal Care and Use Committee (IACUC), and mouse embryos were collected by following the American Veterinary Medical Association (AVMA) guidelines.

## Consent for publication

All authors read and approved the final manuscript.

## Availability of data and materials

The mx-scRNA-seq datasets generated and/or analysed during the current study are available in BioStudies (Acces-sion No. E-MTAB-13181) [https://www.ebi.ac.uk/biostudies/arrayexpress/studies/E-MTAB-13181]. The sample-barcode read matrices and the R scripts for estimating the multiplet probabilities used in this study are available in Github (https://github.com/fnakaki/stealth-multiplet-notebook).

## Authors’ contributions

F.N. conceived the project, build the theoretical frame-work, performed the mx-scRNA-seq, analysed the data, and wrote the manuscript. J.S. conceived the project and wrote the manuscript.

## Acknowledgments

We thank EMBL Heidelberg Genomics Core Facility (GeneCore) and UPF Genomics Core Facility for technical supports, PRBB Animal Facility for animal care, EMBL IT Service for the service of high perfomance computing resources, Heura Cardona and Kerim Anlas for providing cell samples with us, Marco Musy for critical review on mathematics, Xavier Diego, James Cotterell, and Antoni Matyjaszkiewicz for constructive discussion and comments on the paper, Zev Gartner, Lars Steinmetz, Daniel Schraivogel, and Vladimir Benes for invaluable comments on the paper, Qin Zhu for his great insight into demultiplexing algorithms.

## References

[1] Klein AM, Mazutis L, Akartuna I, Tallapragada N, Veres A, Li V, et al. Droplet Barcoding for Single-Cell Transcriptomics Applied to Embryonic Stem Cells. Cell. 2015 May;161(5):1187–1201. Publisher: Elsevier Inc.. 10.1016/j.cell.2015.04.044.

[2] Macosko EZ, Basu A, Satija R, Nemesh J, Shekhar K, Goldman M, et al. Highly Parallel Genome-wide Expression Profiling of Individual Cells Using Nanoliter Droplets. Cell. 2015 May;161(5):1202–1214. Publisher: Elsevier Inc.. 10.1016/j.cell.2015.05.002.

[3] Gierahn TM, Wadsworth MH, Hughes TK, Bryson BD, Butler A, Satija R, et al. Seq-Well: portable, low-cost RNA sequencing of single cells at high throughput. Nature Methods. 2017 Apr;14(4):395–398. 10.1038/nmeth.4179.

[4] Han X, Wang R, Zhou Y, Fei L, Sun H, Lai S, et al. Mapping the Mouse Cell Atlas by Microwell-Seq. Cell. 2018 Feb;172(5):1091–1107.e17. 10.1016/j.cell.2018.02.001.

[5] Rosenberg AB, Roco CM, Muscat RA, Kuchina A, Sample P, Yao Z, et al. Single-cell profiling of the developing mouse brain and spinal cord with split-pool barcoding. Science. 2018 Apr;360(6385):176–182. Publisher: American Association for the Advancement of Science. 10.1126/science.aam8999.

[6] Zheng GXY, Terry JM, Belgrader P, Ryvkin P, Bent ZW, Wilson R, et al. Massively parallel digital transcriptional profiling of single cells. Nature Communications. 2017 Jan;8(1):14049. 10.1038/ncomms14049.

[7] Svensson V, Vento-Tormo R, Teichmann SA. Exponential scaling of single-cell RNA-seq in the past decade. Nature Protocols. 2018 Apr;13(4):599–604. 10.1038/nprot.2017.149.

[8] Haque A, Engel J, Teichmann SA, Lönnberg T. A practical guide to singlecell RNA-sequencing for biomedical research and clinical applications. Genome Medicine. 2017 Dec;9(1):75. 10.1186/s13073-017-0467-4.

[9] Wolock SL, Lopez R, Klein AM. Scrublet: Computational Identification of Cell Doublets in Single-Cell Transcriptomic Data. Cell Systems. 2019 Apr;8(4):281– 291.e9. 10.1016/j.cels.2018.11.005.

[10] McGinnis CS, Murrow LM, Gartner ZJ. DoubletFinder: Doublet Detection in Single-Cell RNA Sequencing Data Using Artificial Nearest Neighbors. Cell Systems. 2019 Apr;8(4):329–337.e4. 10.1016/j.cels.2019.03.003.

[11] DePasquale EAK, Schnell DJ, Van Camp PJ, Valiente-Alandí In, Blaxall BC, Grimes HL, et al. DoubletDecon: Deconvoluting Doublets from Single-Cell RNA-Sequencing Data. Cell Reports. 2019 Nov;29(6):1718–1727.e8. 10.1016/j.celrep.2019.09.082.

[12] Germain PL, Lun A, Macnair W, Robinson MD. Doublet identification in singlecell sequencing data using scDblFinder. F1000Research. 2021 Sep;10:979. 10.12688/f1000research.73600.1.

[13] Stoeckius M, Hafemeister C, Stephenson W, Houck-Loomis B, Chattopadhyay PK, Swerdlow H, et al. Simultaneous epitope and transcriptome measurement in single cells. Nature Methods. 2017 Sep;14(9):865–868. 10.1038/meth.4380.

[14] McGinnis CS, Patterson DM, Winkler J, Conrad DN, Hein MY, Srivastava V, et al. MULTI-seq: sample multiplexing for single-cell RNA sequencing using lipid-tagged indices. Nature Methods. 2019 Jul;16(7):619–626. 10.1038/s41592-019-0433-8.

[15] Gaublomme JT, Li B, McCabe C, Knecht A, Yang Y, Drokhlyansky E, et al. Nuclei multiplexing with barcoded antibodies for single-nucleus genomics. Nature Communications. 2019 Jul;10(1):2907. 10.1038/s41467-019-10756-2.

[16] Shin D, Lee W, Lee JH, Bang D. Multiplexed single-cell RNA-seq via transient barcoding for simultaneous expression profiling of various drug perturbations. Science Advances. 2019 May;5(5):eaav2249. 10.1126/sciadv.aav2249.

[17] Gehring J, Hwee Park J, Chen S, Thomson M, Pachter L. Highly multiplexed single-cell RNA-seq by DNA oligonucleotide tagging of cellular proteins. Nature Biotechnology. 2020 Jan;38(1):35–38. 10.1038/s41587-019-0372-z.

[18] Cheng J, Liao J, Shao X, Lu X, Fan X. Multiplexing Methods for Simultaneous Large-Scale Transcriptomic Profiling of Samples at Single-Cell Resolution. Advanced Science. 2021 Sep;8(17):2101229. 10.1002/advs.202101229.

[19] Kang HM, Subramaniam M, Targ S, Nguyen M, Maliskova L, McCarthy E, et al. Multiplexed droplet single-cell RNA-sequencing using natural genetic variation. Nature Biotechnology. 2018 Jan;36(1):89–94. 10.1038/nbt.4042.

[20] Guo C, Kong W, Kamimoto K, Rivera-Gonzalez GC, Yang X, Kirita Y, et al. Cell-Tag Indexing: genetic barcode-based sample multiplexing for single-cell genomics. Genome Biology. 2019 Dec;20(1):90. 10.1186/s13059-019-1699-y.

[21] Stoeckius M, Zheng S, Houck-Loomis B, Hao S, Yeung BZ, Mauck WM, et al. Cell Hashing with barcoded antibodies enables multiplexing and doublet detection for single cell genomics. Genome Biology. 2018 Dec;19(1):224. 10.1186/s13059-018-1603-1.

[22] Xin H, Lian Q, Jiang Y, Luo J, Wang X, Erb C, et al. GMM-Demux: sample demultiplexing, multiplet detection, experiment planning, and novel cell-type verification in single cell sequencing. Genome Biology. 2020 Dec;21(1):188. 10.1186/s13059-020-02084-2.

[23] Bloom JD. Estimating the frequency of multiplets in single-cell RNA sequencing from cell-mixing experiments. PeerJ. 2018 Sep;6:e5578. 10.7717/peerj.5578.

[24] Hao Y, Hao S, Andersen-Nissen E, Mauck WM, Zheng S, Butler A, et al. Integrated analysis of multimodal single-cell data. Cell. 2021 Jun;184(13):3573– 3587.e29. 10.1016/j.cell.2021.04.048.

[25] Boggy GJ, McElfresh GW, Mahyari E, Ventura AB, Hansen SG, Picker LJ, et al. BFF and cellhashR: analysis tools for accurate demultiplexing of cell hashing data. Bioinformatics. 2022 May;38(10):2791–2801. 10.1093/bioinformatics/btac213.

[26] Klein HU. demuxmix: demultiplexing oligonucleotide-barcoded single-cell RNA sequencing data with regression mixture models. Bioinformatics. 2023 Aug;39(8):btad481. 10.1093/bioinformatics/btad481.

[27] Zhu Q, Conrad DN, Gartner ZJ. deMULTIplex2: robust sample demultiplexing for scRNA-seq. Bioinformatics; 2023. Available from: 10.1101/2023.04.11.536275.

[28] Lareau CA, Ma S, Duarte FM, Buenrostro JD. Inference and effects of barcode multiplets in droplet-based single-cell assays. Nature Communications. 2020 Feb;11(1):866. Number: 1 Publisher: Nature Publishing Group. 10.1038/s41467-020-14667-5.

[29] Datlinger P, Rendeiro AF, Boenke T, Senekowitsch M, Krausgruber T, Barreca D, et al. Ultra-high-throughput single-cell RNA sequencing and perturbation screening with combinatorial fluidic indexing. Nature Methods. 2021 Jun;18(6):635–642. 10.1038/s41592-021-01153-z.

[30] R Core Team. R: A Language and Environment for Statistical Computing. Vienna, Austria: R Foundation for Statistical Computing; 2020. Available from: https://www.R-project.org/.

[31] Nakamura Y, Yamamoto K, He X, Otsuki B, Kim Y, Murao H, et al. Wwp2 is essential for palatogenesis mediated by the interaction between Sox9 and mediator subunit 25. Nature Communications. 2011 Mar;2:251–10. Publisher: Nature Publishing Group. 10.1038/ncomms1242.

[32] George SHL, Gertsenstein M, Vintersten K, Korets-Smith E, Murphy J, Stevens ME, et al. Developmental and adult phenotyping directly from mutant embryonic stem cells. Proceedings of the National Academy of Sciences. 2007 Mar;104(11):4455–4460. 10.1073/pnas.0609277104.

[33] Anlas K, Baillie-Benson P, Arato K, Turner DA, Trivedi V. Gastruloids: Embryonic Organoids from Mouse Embryonic Stem Cells to Study Patterning and Development in Early Mammalian Embryos. In: Ebrahimkhani MR, Hislop J, editors. Programmed Morphogenesis: Methods and Protocols. Methods in Molecular Biology. New York, NY: Springer US; 2021. p. 131–147. Available from: 10.1007/978-1-0716-1174-610.

[34] Griffiths JA, Richard AC, Bach K, Lun ATL, Marioni JC. Detection and removal of barcode swapping in single-cell RNA-seq data. Nature Communications. 2018 Jul;9(1):2667. Number: 1 Publisher: Nature Publishing Group. 10.1038/s41467-018-05083-x.

[35] McCarthy DJ, Campbell KR, Lun ATL, Wills QF. Scater: pre-processing, quality control, normalization and visualization of single-cell RNA-seq data in R. Bioinformatics. 2017 Jan; p. btw777. 10.1093/bioinformatics/btw777.

[36] Lun ATL, McCarthy DJ, Marioni JC. A step-by-step workflow for low-level analysis of single-cell RNA-seq data with Bioconductor. F1000Research; 2016. 5:2122. Type: article. Available from: https://f1000research.com/articles/5-2122.

[37] Toedling J, Sklyar O, Huber W. Ringo – an R/Bioconductor package for analyzing ChIP-chip readouts. BMC Bioinformatics. 2007 Jun;8(1):221. 10.1186/1471-2105-8-221.

