## Supplementary Material for "Probability of stealth multiplets in sample-multiplexing for droplet-based single-cell analysis"

### Modelling a multiplet probability by the Poisson distribution

The Poisson distribution has been applied to account for the frequency of cell encapsulation in the droplet-based single-cell analysis [1–4]. Indeed, it meets the assumptions of the distribution that are A) Random and B) Low frequency. The distribution is expressed as,

$$P(k) = \frac{\lambda^k e^{-\lambda}}{k!} \qquad (0 < \lambda < 1)$$

where $\lambda$ is the average count of the event, representing the average cell number per droplet. Because cells are loaded with low concentration to minimise the multiplet generation, $\lambda$ must be $0 < \lambda < 1$. $k$ is the number of events, which means the number of cells coming into a single droplet. The probability of empty droplets , singlets, doublets, and $k$-ets (with or without beads) can be represented as $P(0)$, $P(1)$, $P(2)$, and $P(k)$, respectively. Therefore, the probability of multiplets is,

$$P(k \geq 2) = 1 - P(0) - P(1) = 1 - e^{-\lambda} - \lambda e^{-\lambda}$$

**Supplementary Table 1** showed the probabilities of singlets, doublets, and triplets... based on the Poisson distribution. In this case, 0-et means an empty droplet, and these probabilities are among total droplets of which the number is usually unknown. Instead, what can be observed is the total number of the cell-droplets ($= k \geq 1$), and the multiplet ratio, the fraction of multiplets among the cell-droplets can be expressed as,

$$(multiplet\ ratio) = \frac{P(k \geq 2)}{P(k \geq 1)} = 1 - \frac{P(1)}{1 - P(0)} = 1 - \frac{\lambda e^{-\lambda}}{1 - e^{-\lambda}} \tag{1}$$

Also, the fraction of the singlet, doublet, triplet,... under the corresponding $\lambda$ is shown in **Supplementary Table 2**. More than 80% was the singlet at around $\lambda \leq 0.4$. Indeed, the Chromium Controller (10X Genomics), which is one of the common microfluidics platforms for the droplet-based scRNA-seq, originally recommended the target cell number should be less than 10,000 cells, yielding 8% of multiplet among all cell-droplet, which corresponds to $\lambda = 0.166$. Loading cells exceeding this upper limit is regarded as "super-loading" ($\lambda > 0.166$) [5]. Indeed, at $\lambda = 0.5$, around 4% of multiplets are quartets or more, according to **Supplementary Table 2**. The relation of input cell numbers and $\lambda$ in the Chromium Controller (10X Genomics) are further discussed in the following section.

| $k$-et | $\lambda = 0.1$ | $\lambda = 0.2$ | $\lambda = 0.3$ | $\lambda = 0.4$ | $\lambda = 0.5$ |
|---|---|---|---|---|---|
| 0 | 0.9048374 | 0.8187308 | 0.7408182 | 0.6703200 | 0.6065307 |
| 1 | 0.0904837 | 0.1637462 | 0.2222455 | 0.2681280 | 0.3032653 |
| 2 | 0.0045242 | 0.0163746 | 0.0333368 | 0.0536256 | 0.0758163 |
| 3 | 0.0001508 | 0.0010916 | 0.0033337 | 0.0071501 | 0.0126361 |
| 4 | 0.0000038 | 0.0000546 | 0.0002500 | 0.0007150 | 0.0015795 |
| 5 | 0.0000001 | 0.0000022 | 0.0000150 | 0.0000572 | 0.0001580 |
| 6 | 0.0000000 | 0.0000001 | 0.0000008 | 0.0000038 | 0.0000132 |
| 7 | 0.0000000 | 0.0000000 | 0.0000000 | 0.0000002 | 0.0000009 |
| 8 | 0.0000000 | 0.0000000 | 0.0000000 | 0.0000000 | 0.0000001 |
| 9 | 0.0000000 | 0.0000000 | 0.0000000 | 0.0000000 | 0.0000000 |
| 10 | 0.0000000 | 0.0000000 | 0.0000000 | 0.0000000 | 0.0000000 |

**Supplementary Table 1:** The theoretical probability of droplets containing $k$ cells

| $k$-et | $\lambda = 0.1$ | $\lambda = 0.2$ | $\lambda = 0.3$ | $\lambda = 0.4$ | $\lambda = 0.5$ |
|---|---|---|---|---|---|
| 1 | 0.9508332 | 0.9033311 | 0.8574888 | 0.8132979 | 0.7707470 |
| 2 | 0.0475417 | 0.0903331 | 0.1286233 | 0.1626596 | 0.1926868 |
| 3 | 0.0015847 | 0.0060222 | 0.0128623 | 0.0216879 | 0.0321145 |
| 4 | 0.0000396 | 0.0003011 | 0.0009647 | 0.0021688 | 0.0040143 |
| 5 | 0.0000008 | 0.0000120 | 0.0000579 | 0.0001735 | 0.0004014 |
| 6 | 0.0000000 | 0.0000004 | 0.0000029 | 0.0000116 | 0.0000335 |
| 7 | 0.0000000 | 0.0000000 | 0.0000001 | 0.0000007 | 0.0000024 |
| 8 | 0.0000000 | 0.0000000 | 0.0000000 | 0.0000000 | 0.0000001 |
| 9 | 0.0000000 | 0.0000000 | 0.0000000 | 0.0000000 | 0.0000000 |
| 10 | 0.0000000 | 0.0000000 | 0.0000000 | 0.0000000 | 0.0000000 |

**Supplementary Table 2:** The theoretical ratio of the droplets containing $k$ cells among all cell-droplets

### $\lambda$ in the Chromium Controller (10X Genomics)

Although the Poisson distribution is suitable to describe the probability of cell-droplets, the major drawback in practice is that we cannot see the total number of droplets, and hence, cannot know the key parameter, $\lambda$. However, we can estimate it from the multiplets ratios of the system we use. Here we estimate $\lambda$ in the Chromium Controller (10X Genomics). According to 10X Genomics, the multiplet rates with the Chromium Controller are shown in **Supplementary Figure 2A**. Here, $(\textit{mulitplet ratio}) = 8 \times 10^{-6} N$, where $N$ is the target cell number for analysis. Note that actual loading cell numbers are 165% of the target cell count to recover (from CG000315 Rev B Chromium Next GEM Single Cell 3' Reagent Kits v3.1 (Dual Index) USER GUIDE). These multiplet rates can also be expressed with the Poisson distribution (**Equation 1**). Therefore,

$$\begin{aligned} 8 \times 10^{-6} N &= 1 - \frac{\lambda e^{-\lambda}}{1 - e^{-\lambda}} \\ N &= \left(1 - \frac{\lambda e^{-\lambda}}{1 - e^{-\lambda}}\right) \times 1.25 \times 10^{5} \end{aligned} \tag{2}$$

This relation between $\lambda$ and the target cell counts is shown in **Supplementary Figure 2B**. Because this equation cannot be analytically solved for $\lambda$, a series of the estimated target cell number from $\lambda$ by the equation 2, were plotted, followed by approximation by linear regression (red line, **Supplementary Figure 2B**). The approximated $\lambda$ for each target cell number is shown in **Supplementary Table3**. In the case of the two datasets in this paper, we loaded cells to target 4,000 cells, corresponding to $\lambda = 0.060$, in theory.

$$\begin{aligned} N &\sim 56800\lambda + 570 \\ \lambda &\approx N/56800 - 0.01 \end{aligned}$$

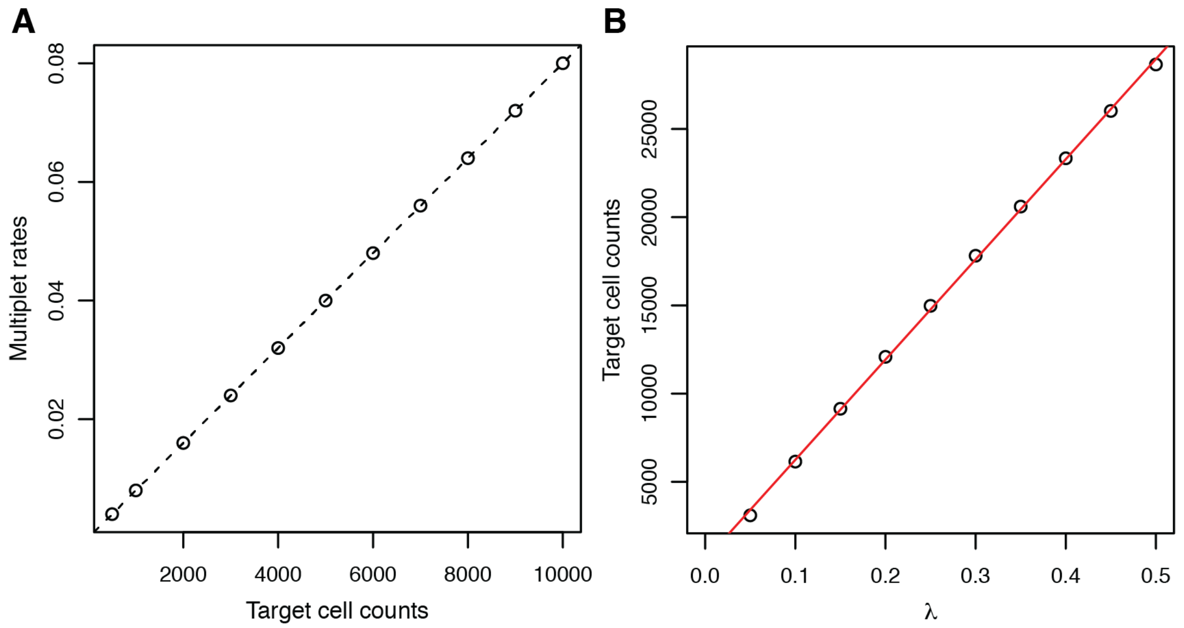


**Supplementary Figure 2:** (A) The relation between target cell counts and multiplet rates according to the Chromium Next GEM Single Cell 3' Reagent Kits v3.1 USER GUIDE (CG000315 Rev B). (B) The estimated relation between $\lambda$ and target cell count. The points were plotted based on **Equation 2**, and the red line was drawn by linear regression for approximation.

| $N$ | Multiplet ratio | $\lambda$ |
|---|---|---|
| 2000 | 0.016 | 0.025 |
| 4000 | 0.032 | 0.060 |
| 6000 | 0.048 | 0.096 |
| 8000 | 0.064 | 0.131 |
| 10000 | 0.080 | 0.166 |
| 12000 | 0.096 | 0.201 |
| 14000 | 0.112 | 0.236 |
| 16000 | 0.128 | 0.272 |
| 18000 | 0.144 | 0.307 |
| 20000 | 0.160 | 0.342 |
| 22000 | 0.176 | 0.377 |
| 24000 | 0.192 | 0.413 |

**Supplementary Table 3:** Estimated $\lambda$ from the multiplet ratio in the Chromium Controller (10x Genomics)

### References

1. Zheng, G. X. Y. *et al.* Massively parallel digital transcriptional profiling of single cells. en. *Nature Communications* **8,** 14049. ISSN: 2041-1723. `https://www.nature.com/articles/ncomms14049` (2023) (Jan. 2017).
2. McGinnis, C. S., Murrow, L. M. & Gartner, Z. J. DoubletFinder: Doublet Detection in Single-Cell RNA Sequencing Data Using Artificial Nearest Neighbors. en. *Cell Systems* **8,** 329–337.e4. ISSN: 24054712. `https://linkinghub.elsevier.com/retrieve/pii/S2405471219300730` (2022) (Apr. 2019).
3. Bloom, J. D. Estimating the frequency of multiplets in single-cell RNA sequencing from cell-mixing experiments. en. *PeerJ* **6,** e5578. ISSN: 2167-8359. `https://peerj.com/articles/5578` (2022) (Sept. 2018).
4. Xin, H. *et al.* GMM-Demux: sample demultiplexing, multiplet detection, experiment planning, and novel cell-type verification in single cell sequencing. en. *Genome Biology* **21,** 188. ISSN: 1474-760X. `https://genomebiology.biomedcentral.com/articles/10.1186/s13059-020-02084-2` (2022) (Dec. 2020).
5. Stoeckius, M. *et al.* Cell Hashing with barcoded antibodies enables multiplexing and doublet detection for single cell genomics. *Genome Biology* **19,** 224. ISSN: 1474-760X. `https://doi.org/10.1186/s13059-018-1603-1` (2023) (Dec. 2018).
